# Molecular origins of heterogeneous aging and spatial organization in RNA condensates

**DOI:** 10.64898/2026.08.27.747560

**Authors:** Dibyajyoti Mohanta, Heyang Zhang, D. Thirumalai, Hung T. Nguyen

## Abstract

The molecular origins of aging of biomolecular condensates, which play a central role in cellular organization, is poorly understood. Here, we use coarse-grained molecular simulations to investigate how RNA sequence and chain connectivity govern condensate aging over extended timescales. Condensates formed by CAG-repeat RNA undergo pronounced aging characterized by progressive dynamical slowing, loss of ergodicity, and the emergence of two distinct relaxation timescales. Aging proceeds heterogeneously in space, giving rise to a dynamically arrested, solid-like core surrounded by a more fluid shell. We demonstrate that aging is driven by sequence-encoded base pairing that favors RNA expansion, alignment and the formation of a dense interchain interaction network. These structural changes lead to increased topological entanglement, stabilizing long-lived conformations and reinforcing dynamic arrest in the condensate interior. Strikingly, a scrambled sequence with identical composition remains largely liquid-like. Our results establish RNA sequence patterning as a key determinant not only of phase separation but also of condensate aging and spatial organization. These findings provide a molecular framework for understanding the persistence and solidification of repeat RNA assemblies observed in diseases and suggest general physical principles by which entangled polymer networks drive aging in biomolecular condensates.

## I. INTRODUCTION

Cells organize their biochemical reactions through both membrane-bound organelles and a diverse array of membraneless biomolecular condensates that form via phase separation.^1–4^ While the initial stages of condensate formation have been extensively characterized, much less is known about the long-time evolution of their properties, broadly referred to as condensate aging.^4–8^ Aging describes the gradual transition of condensates from fluid-like droplets to viscoelastic, gel-like, or even solid-like states over timescales ranging from minutes to months.^9,10^ This process results in progressive slowing of molecular diffusion, loss of the ability of droplets to fuse, and the emergence of conformational heterogeneity, as observed in glasses.^11,12^ The physical and biological implications of condensate aging are profound: while modest increases in condensate viscosity may stabilize functional assemblies, excess hardening or irreversible aggregation is implicated in neurodegenerative disorders (ALS and FTD) and cancer.^13–17^ In this sense, condensate aging bridges the continuum between functional, dynamic cellular organization and pathological, arrested states.^13,18,19^

One hallmark of aging is the emergence of internal structural and compositional heterogeneity.^20^ Rather than remaining homogeneous liquids, aging condensates often evolve into core-shell architectures.^10,21–24^ Experimental studies have revealed that subregions inside condensates progressively solidify due to irreversible cross-linking, while the rest of the regions retains residual fluidity that allows limited molecular exchange with the surroundings.^8–10,25,26^ This “phase-within-phase” organization is a natural consequence of kinetic arrest and differential molecular mobility. Similar multiphase architectures have also been observed in RNA-protein condensates, where distinct local compositions arise from preferential partitioning of RNAs and proteins with different valence or charge patterns.^27–32^

The molecular origins of condensate aging are complex and multifaceted. Several mechanisms have been proposed, including liquid-to-solid transitions driven by structural rearrangements, covalent or noncovalent cross-linking, and chemical modifications that alter molecular interactions.^10,16,24,26,33–40^ Fluorescense recovery after photobleaching (FRAP) and particle tracking microrheology have revealed a spectrum of material transitions from purely viscous liquids to viscoelastic gels and amorphous solids, underscoring the diversity of physical states accessible to condensates.^6,8,41^ Computations have also played a pivotal role in illuminating the microscopic determinants of condensate formation.^42–50^ However, the molecular mechanisms underlying condensate aging are still elusive.^4,51^ All-atom molecular dynamics (MD) simulations can reveal detailed interaction motifs and hydration structures but are limited to microsecond timescales and nanometer length scales.^10,52,53^ Coarsegrained (CG) models simplify molecular representations and effective interactions, allowing simulations of droplet coalescence and viscoelastic relaxation.^54,55^ Importantly, previous CG simulations, mostly in the context of intrinsically disordered proteins, have already reproduced several hallmarks of condensate aging, including the emergence of dense core-shell organization, orientational ordering, and slow internal dynamics.^10,35,36,56,57^ However, a fundamental question remains: can condensate aging arise spontaneously from sequence-encoded molecular interactions, and if so, what are the microscopic mechanisms that govern its temporal evolution and spatial heterogeneity?

Here, we present CG simulations that capture the aging behavior of CAG-repeat RNA condensates, implicated in neurodegeneration diseases,^58^ without altering the interaction parameters in an *ad hoc* manner. We show that dynamical slowing down and structural reorganization inferred experimentally^13,59^ occur spontaneously in our simulations, thereby demonstrating that aging is intrinsic to the sequence-dependent interactions and chain connectivity. Strikingly, we uncover a previously uncharacterized spatiotemporal dependence of condensate aging: distinct regions within a single droplet exhibit markedly different dynamical and structural evolution, revealing that aging is inherently heterogeneous. We find that heterogeneity originates primarily due to differential base-pairing interactions, which drive the formation of dense, slowly relaxing RNA networks in the condensate core while maintaining a more dynamic and fluid outer shell. In contrast, a scrambled RNA sequence that lacks complementary base-pairing exhibits minimal aging behavior. Together, these findings establish a direct molecular link between the prevalence of homotypic RNA interactions, irreversible aggregation, and the gradual emergence of arrested dynamics within RNA condensates.

## II. RESULTS

### A. Aging in Condensates of CAG Repeats

We begin by examining how the (CAG)_31_ repeat condensate evolves over time (Fig. 1). Due to the continuous exchange between the dilute and condensed phases, we chose to only track molecules that consistently stay in the droplet over the simulation time scale.

**FIG. 1.**
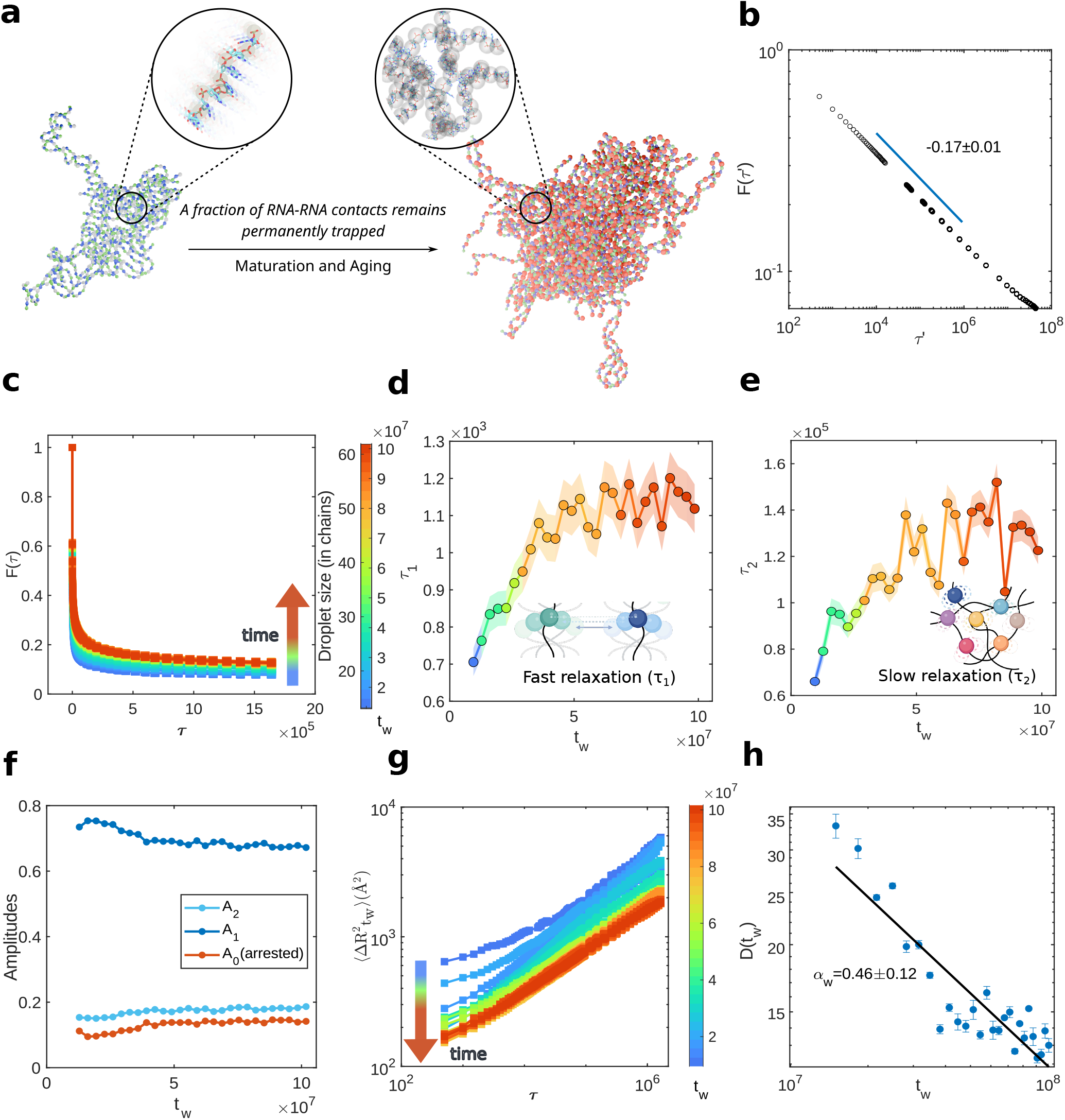
Aging dynamics of CAG-repeat RNA condensates. (a) Representative snapshots showing evolution of condensate from early age (*t*_*w*_ = 10^7^ time steps, left; blue inset) to mature age (*t*_*w*_ = 10^8^ timesteps, right; red inset). Colored spheres represent RNA nucleotides. Early-age inset: RNA chains exhibit rapid, large-amplitude cage rattling motion, indicated by extended motion trails of gray beads. Mature-age inset: a fraction of molecules becomes kinetically trapped within a confined local environment, with minimal net displacement represented by restricted motion trails of red beads. (b) Time-averaged pairwise overlap function *F*(*τ* ′) over the whole trajectory of 30 chains which are consistently in the droplet. (c) Age-dependent overlap function 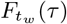 as the droplet ages. The color gradient (blue → red) shows the age progression of the system. The upward shift of 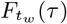 curves with *t*_*w*_ reflects increasing correlation and dynamic slowdown. (d) and (e) Relaxation time scales *τ*_1_ and *τ*_2_ (Eq. 6) increase as the droplet ages, indicating the system undergoes aging with time. We extract *τ*_1_ (fast, rattling motion) and *τ*_2_ (slow, collective rearrangements) from a bi-exponential fit of 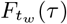. Note that *τ*_1_ and *τ*_2_ differ by two orders of magnitude. (f) Amplitudes from bi-exponential fit with *A*_0_ (arrested fraction) increasing modestly. *A*_1_ and *A*_2_ are fractions of fast and slow relaxation, respectively. (g) Pairwise mean square displacement 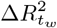 between chains in the droplet for different ages *t*_*w*_ . (h) Average generalized diffusion coefficient *D*(*t*_*w*_) decreases as *t*_*w*_ increases. Error bars represent the standard deviation within the time window *t*_*w*_ . Power-law fit with exponent *α*_*w*_ = 0.46 *±* 0.12, where the uncertainty represents the 95% confidence interval obtained from linear regression analysis.

To quantify internal structural rearrangements, we consider all the non-adjacent nucleotide pairs *i* and *j* within the condensate, including both intra- and inter-chain interactions. For each pair, we define the instantaneous separation distance as:

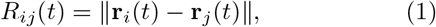

where **r**_*i*_(*t*) denotes the position of nucleotide *i* at time *t*. The displacement of this pairwise distance over a time interval [*t*_1_, *t*_2_] is then given by:

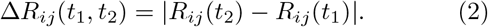

We then characterize the structural memory of the RNA condensate using the overlap function *F* (*τ*′), constructed from Δ*R*_*ij*_(*t*_1_, *t*_2_) and evaluated over the lifetime of the droplet. This metric has been used to probe protein folding dynamics^60^ as well as non-equilibrium relaxation and dynamical arrest in soft matter systems.^61,62^ The overlap function is defined as:

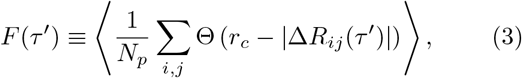

where Θ is the Heaviside step function, contributing only when the relative displacement is smaller than a cut-off distance *r*_*c*_, chosen here as the Lennard–Jones diameter of the nucleotide (*σ* = 5.9 ^Å^). The normalization factor *N*_*p*_ is the number of pairs (*i, j*) satisfying |*i* −*j*| *>* 1, thereby excluding adjacent nucleotides in the same molecule. *F* (*τ*′) quantifies the fraction of pairs that retain their relative positions within the cutoff *r*_*c*_ after a time delay *τ*′. In simple liquids, it typically decays exponentially from unity to zero at long times, reflecting complete structural relaxation and ergodic behavior. In contrast, for the CAG-repeat droplet, *F* (*τ*′) does not decay to 0 even at the longest accessible timescales (Figs. 1b, S2), indicating persistent structural memory and incomplete relaxation within the droplet. Notably, the decay follows a power-law form,

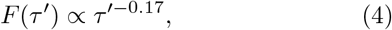

which is reminiscent of the stress relaxation in crosslinked polymer networks.^63–65^ The power law relaxation would be consistent with slow, glassy dynamics with long-lived correlations. This behavior motivates a more detailed analysis of aging by resolving the dynamics across successive temporal windows.

To this end, we partition the trajectory into *n* = 29 non-overlapping sequential windows, after excluding the initial period required for droplet formation. Within each window, the overlap function is calculated as:

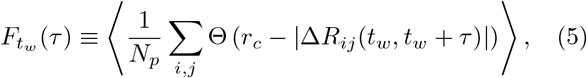

where *t*_*w*_ denotes the starting time of window *w*, and *τ* is the observation time within that window. Fig. 1c shows the decay of 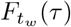 as a function of *τ* for successive windows, with color indicating increasing condensate age (*t*_*w*_, blue → red). Within each window, 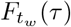 decays from unity to approximately ∼0.1, indicating partial decorrelation of nucleotide pairs while retaining significant structural memory. More intriguingly, as the waiting time *t*_*w*_ increases, the decay of 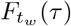 becomes progressively slower, manifested as an upward shift of the curves. This systematic slowdown is a hallmark of aging dynamics, where relaxation times increase with system age and correlations become increasingly long-lived.^16,66^

A bi-exponential function provides the best fit to 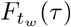 within each time windows (see Fig. S3):

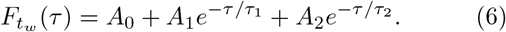

This decomposition reveals two distinct relaxation timescales: a *fast* mode *τ*_1_ and a *slow* mode *τ*_2_, characteristic of heterogeneous relaxation in soft and glassy materials.^67,68^ The coefficient *A*_0_ corresponds to a non-relaxing (arrested) fraction, associated with contacts whose relative positions remain stable over the entire observation window, analogous to the non-ergodic parameter in structural glasses.^69,70^ *A*_1_ and *A*_2_ represent the population fractions of the fast and slow relaxation modes, respectively, subject to the normalization constraint ∑_*k*=0,1,2_ *A*_*k*_ = 1. The fit parameters for all aging windows are shown in Fig. S4, with excellent agreement in all cases (*R*^2^ *>* 0.95).

Both relaxation timescales, *τ*_1_ and *τ*_2_, exhibit similar aging behavior: they increase rapidly with the waiting time *t*_*w*_ before saturating at longer ages (Figs. 1d, e). At the same time, the fraction of the fast relaxation mode (*A*_1_) decreases steadily, while both the slow mode (*A*_2_) and arrested fraction (*A*_0_) increase modestly with age (Fig. 1f). This evolution is consistent with the emergence of slow dynamics and progressive dynamical arrest in aging polymer networks.^71,72^ The coexistence of two relaxation modes and their evolution with age underlies the observed power-law relaxation in our simulations.^64,73^

To further quantify aging-induced changes in structural relaxation, we analyze the pairwise mean square displacement (pMSD), 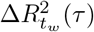,^57^ defined over a lag time *τ* = *t*_2_ − *t*_1_ as:

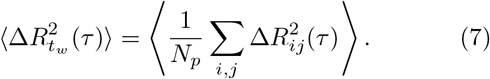

The pMSD captures the time-averaged relative displacement between all pairs as a function of *τ* at a given age *t*_*w*_. Within each window, pMSD increases with the observation time *τ* (Fig. 1g), but its overall magnitude decreases systematically with increasing age, indicating progressive dynamical slowing consistent with the overlap function analysis. The pMSD follows a power-law scaling, 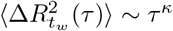, where the exponent *κ* characterizes the nature of the dynamics: normal diffusive (*κ* = 1), subdiffusive (*κ <* 1) or superdiffusive (*κ >* 1). We observe subdiffusion across all ages, with *κ* ∼ 0.30 − 0.40 (Fig. S5). Accounting explicitly for aging effects, we express the pMSD as:

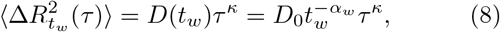

where *α*_*w*_ = 0.46 *±* 0.12 (with *κ* fixed at 0.35) quantifies the age-dependent slowdown of diffusion (Fig. 1h). Such age-induced subdiffusive dynamics have been widely reported in protein and RNA-protein condensates undergoing disorder-order transitions during aging.^6,8,16,36^

### B. Aging is Spatially Heterogeneous

The emergence of two evolving timescales in the overlap function (Eq. 3) is suggestive of heterogeneous relaxation dynamics. Motivated by experiments showing that condensates often exhibit multi-layered organization,^74^ we wonder whether the emergence of the two timescales is a consequence of a heterogeneous environment. Thus, we examine position-dependent dynamics by computing the shell-resolved mean square displacement (MSD). Specifically, nucleotides were partitioned into three concentric spherical shells (core, inner and outer shells), each with thickness Δ*r*_shell_ = 5.0 nm, radiating outward from the droplet center of mass (COM) (Fig. 2a). The shell-wise MSD is defined as:

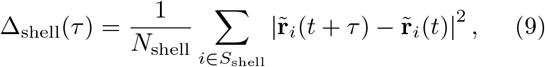

where 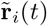 is the position of the *i*-th nucleotide relative to the droplet COM. *N*_shell_ is the number of nucleotides whose radial distances from the COM is within the shell interval Δ*r*_shell_ ∈ *S*_shell_[*R, R* + Δ*r*_shell_] at both the initial time *t* and the later time *t* + *τ* . The shell-specific dynamics *t*_*w*_ are characterized by a generalized diffusion coefficient through:

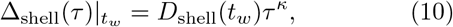

with the subdiffusive exponent fixed at *κ* = 0.35 for all shells (Fig. S6). Remarkably, we find that the nucleotide motion in the core is significantly slower compared with those in the outer regions. The effective diffusion coefficients *D*(*t*_*w*_) exhibits a clear spatial gradient, increasing monotonically from the core toward the condensate surface (Fig. 2b). Importantly, aging effects are most pronounced in the core: the diffusion in the core decays most rapidly with age, with the largest aging exponent *α*_*w*,core_ = 0.53 *±* 0.11, compared to *α*_*w*,Inner shell_ = 0.49 *±* 0.12 in the inner shell and *α*_*w*,Outer shell_ = 0.26 *±* 0.15 near the surface (Fig. 2b). Similar behavior is observed if we instead partition the condensate into four or two shells (Figs. S12 and S13). These results demonstrate that condensate dynamics depend both on the spatial location and droplet age, providing a dynamical basis for the core-shell architectures observed in many biomolecular condensates.^27,75,76^

**FIG. 2.**
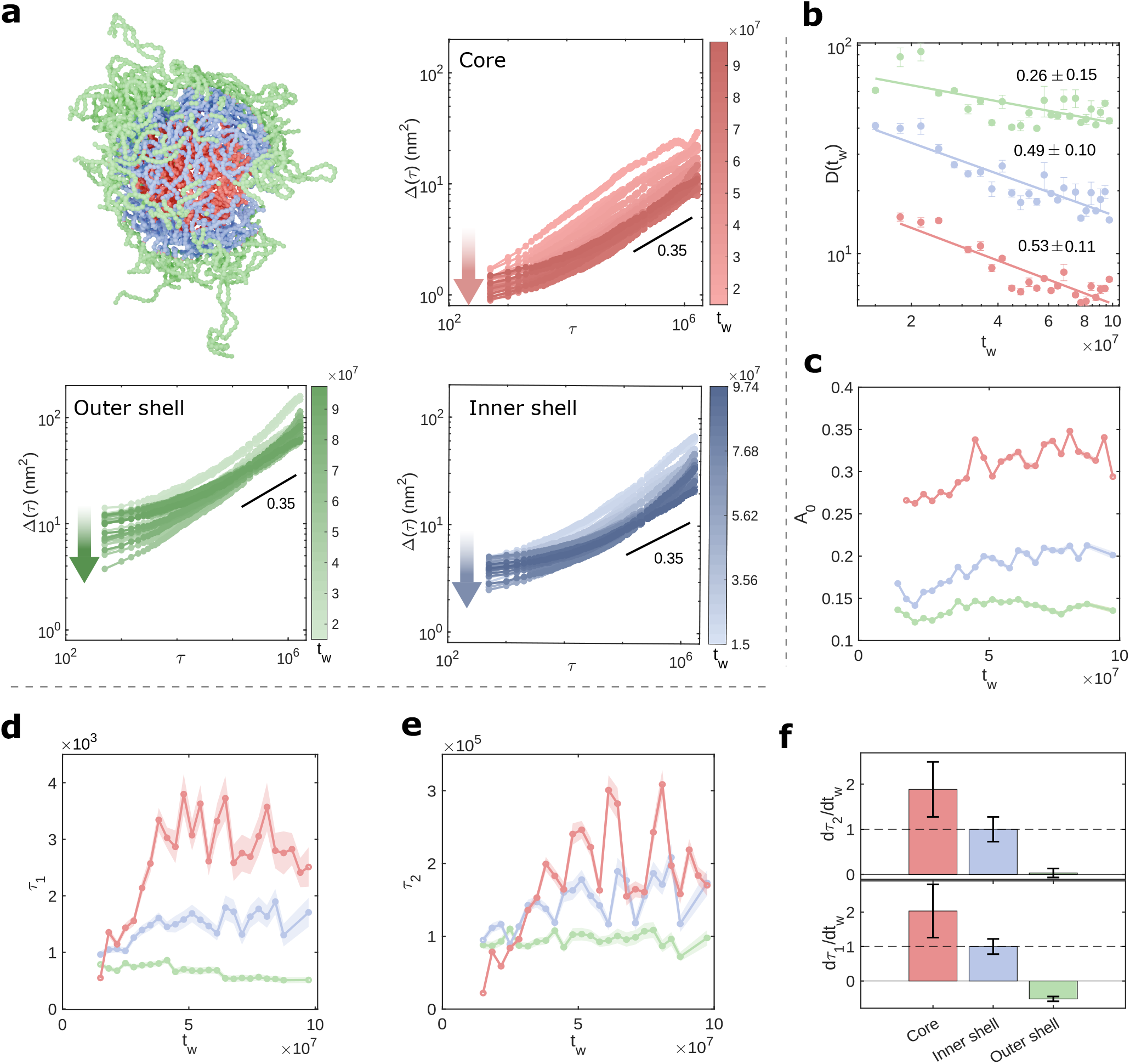
Shell-wise aging in CAG repeat condensate. (a) Illustration of the concentric spherical shells for condensates with thickness Δ*r*_shell_ = 5.0 nm from the droplet’s center-of-mass: core (red), inner shell (blue), and outer shell (green). Although the snapshot contains a few chain ends dangling from the surface, the green region includes only segments within the droplet radius of ≈15 nm. Shell-wise mean squared displacement (MSD) 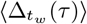 at varying ages *t*_*w*_ (intensity gradients from light to dark represent age progression). The downward trend in MSD curves with increasing *t*_*w*_ shows dynamic slowing in all shells, but with spatial variance: core exhibits the lowest MSD (most restricted), gradually increasing toward the outer shell (greater molecular fluidity). (b) Shell-wise variation of diffusion inside the droplet *D*(*t*_*w*_) as it ages (log-log plot). Error bars represent the standard deviation in *D*(*t*_*w*_) within each time window *t*_*w*_ . The aging exponent *α* is extracted for each shell from the power-law fitting, with uncertainties representing 95% confidence interval. (c) Variation of the arrested fraction *A*_0_ with condensate age *t*_*w*_ (from bi-exponential fit of the shell-wise 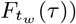. The core exhibits the highest *A*_0_ values and decreases toward outer shells. In all shells, *A*_0_ increases with age, reflecting progressive freezing of contacts over time. (d-e) Fast and slow relaxation timescales are highest in core, decreasing outward. (f) Temporal aging rates of relaxation timescales, *dτ*_1_*/dt*_*w*_ and *dτ*_2_*/dt*_*w*_ with *t*_*w*_ (relative to the rate of the inner shell). Both rates exhibit maximum magnitude at the core, decreasing linearly toward the outer shell.

Given the pronounced spatiotemporal heterogeneity, the two relaxation timescales *τ*_1_ and *τ*_2_ identified for the whole droplet likely reflect a superposition of location-specific relaxation processes. To test this idea, we repeat the bi-exponential decomposition for each shell separately (Figs. S7, S8, and S9). We find that the arrested fraction *A*_0_ has the largest value in the core, reaching ∼0.30, suggesting that nearly 30% of the contacts remain persistent over the observation window (Fig. 2c). In contrast, the inner and outer shells are progressively more deformed, with *A*_0,inner_ ≈ 0.20 and *A*_0,outer_ ≈ 0.13, respectively. Notably, the non-relaxing fraction increases slightly with age at all locations, consistent with the behavior observed for the whole droplet (Fig. 1f). In addition, both relaxation times are largest in the core and decrease gradually toward the surface (Figs. 2d and e), indicating that nucleotides in the core require approximately six times longer to relax than those near the periphery. Correspondingly, the population of the fast mode (*A*_1_) is the smallest in the core and increases toward the outer shell, consistent with increasingly rapid segmental motion near the surface (Fig. S10). Beyond the spatial differences in the relaxation times, the rates at which these timescales evolve with age (*dτ*_1_*/dt*_*w*_ and *dτ*_2_*/dt*_*w*_) are also maximal in the core and decay sharply toward the periphery (Fig. 2f).

Our shell-resolved analyses establish that the dynamics within CAG-repeat RNA condensates has two defining features. First, molecular motion progressively slows as the condensate ages.^16,27,76^ This is a hallmark of aging that has been reported across a wide range of soft and biological materials.^12,77,78^ Second, the dynamics strongly depend on spatial location within the droplet. The interplay of these two effects produces spatial-dependent aging rates, with the core undergoing markedly accelerated aging relative to other regions. If such dynamical differences persist over extended timescales, the core is expected to undergo a transition to a solid-like, dynamically arrested state, while the periphery may retain liquid-like behavior due to its slower aging kinetics.

### C. Conformational Heterogeneity in RNA Condensate

The emergence of a core-shell architecture in RNA condensates likely reflects the different local environment experienced by RNA chains in the condensate interior compared with those at the periphery. Such differences, in turn, influence both RNA conformation and dynamics. To elucidate the molecular origins of this spatial organization, we examine RNA conformations and structures across the condensate volume.

We find that RNA progressively expand when moving from the droplet surface toward the core (Fig. 3a). Chains near the core exhibit significantly larger radii of gyration (*R*_*g*_ ≈ 7.0–7.5 nm) than those in the outer shells (*R*_*g*_ ≈ 5.5–6.0 nm), consistent with our earlier finding and other simulations of low-complexity protein condensates.^49,79–81^ Importantly, as the condensate ages, the rate of chain expansion *d*⟨*R*_*g*_⟩ */dt*_*w*_ is highest in the core, whereas chains in the inner and outer shells expand more slowly, maintaining relatively stable conformations over the aging timescale (Fig. 3b). To further characterize chain conformations across the radial shells, we extract the Flory exponent *ν* from the segment-distance scaling relation ⟨*R*_|*i*−*j*|_⟩ ∼ *b*|*i* − *j*|^*ν*^ (Fig. S20), where *R*_|*i*−*j*|_ is the average spatial distance between nucleotides *i* and *j, b* is the effective Kuhn length, and |*i* − *j*| denotes the contour separation.^82^ The Flory exponent reflects the effective solvent quality experienced by RNA: *ν* = 0.5 corresponds to ideal-chain behavior, *ν* ≈ 0.588 to swollen conformations in good solvent, and *ν* ≈ 0.33 to compact globular states in poor solvent. Within the core, RNA segments exhibit high Flory exponents (*ν >* 0.5), indicative of extended conformations (Fig. 3c). In contrast, chains in the inner and outer shells display progressively smaller exponents (*ν* ≈ 0.45 and 0.40, respectively), reflecting a shift toward more compact, poor-solvent behavior. Thus, the accelerated dynamic arrest observed in the core (Fig. 2) correlates strongly with enhanced RNA expansion in this region.

**FIG. 3.**
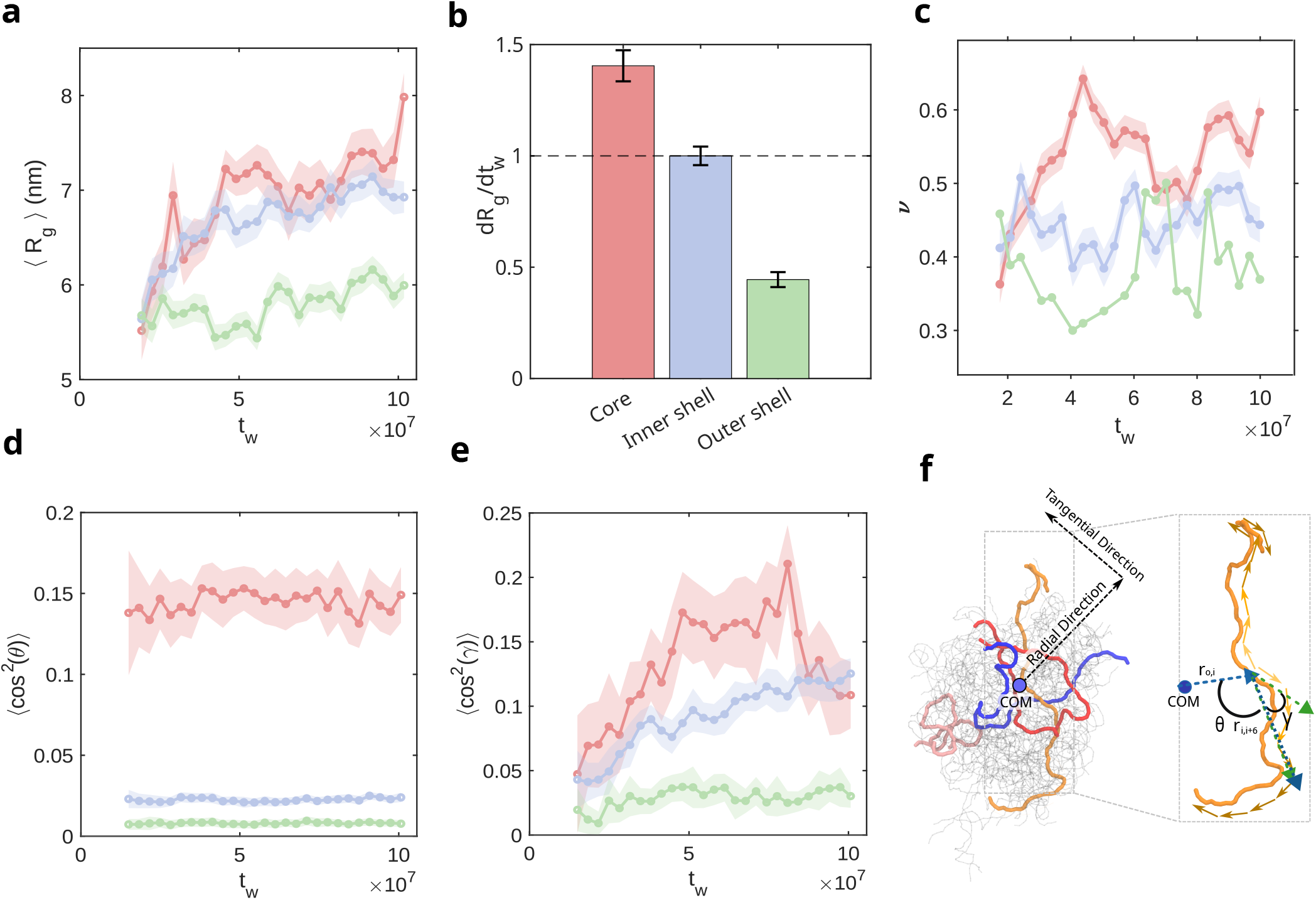
Conformational and orientational heterogeneity in CAG-repeat condensate. (a) Shell-wise radius of gyration ⟨*R*_*g*_⟩ as the condensate ages. Core chains are most extended, while inner and outer shell chains have progressively smaller sizes. (b) The relative rates of *R*_*g*_ increase showing that temporal changes in chain size are largest in the core and progressively decrease toward the periphery. (c) Flory exponent ν, extracted from the scaling relation ⟨*R*_|*i*−*j*|_⟩ ∼ *b* |*i* −*j*| ^*ν*^, as a function of condensate age for different shells. The values in the core are high (ν ≈ 0.5 − 0.6), indicating swollen or extended chain conformations, while surface chains exhibit lower exponents (ν ≈ 0.3 ∼ 0.4), consistent with more compact conformations. (d) Average square cosine of the angle between chain segment vectors 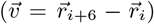 and radial vector 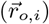 from the droplet center-of-mass (COM) to bead *i* (illustrated in f) as the condensate ages. Values for the outer segments approach 0, indicating near tangential alignment, while core segments exhibit slight orientation toward the COM. (e) Average square cosine of the angle (*γ*) between the *i*-th and (*i* + 6)-th bond vectors within chain segments (illustrated in f). Values for surface segments approach 0, indicating high chain flexibility, while core segments show high persistent alignment and extended conformations. (f) Schematic representation of chain orientations within droplet. Shaded regions in all panels represent standard deviations.

The expansion of chains near the core suggests that their orientational organization may also depend on radial location. To test this hypothesis, we compute the cosine of the angle cos 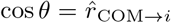 between the radial unit vector from the droplet COM to nucleotide *i* and the unit vector along the chain segment from nucleotide *i* to *i* + 6 (illustrated in Fig. 3f). The squared quantity cos^2^ *θ* quantifies radial alignment: cos^2^ *θ* = 1⟩ indicates perfect radial alignment, whereas cos^2^ *θ* = 0 indicates tangential orientation. We find that segments in the inner and outer shells exhibit small values ⟨cos^2^ *θ*⟩ ≈ 0 (Fig. 3d), indicating a strong preference for tangential alignment around the droplet. In contrast, those in the core display a modest but significant radial bias, with ⟨cos^2^ *θ*⟩ ≈ 0.15 (corresponding to average angles of ∼ 67^*◦*^° or ∼ 113^*◦*^°). This partial radial alignment likely accompanies chain expansion and may enhance inter-chain interlocking and entanglement, thereby progressively hindering relaxation as the condensate ages.^83^

To further quantify orientational persistence, we calculate the cosine of the angle *γ* between bond vectors separated by 6 nucleotides along the chain (Fig. 3f). Larger values of cos^2^ *γ* indicate greater orientational correlation and increased chain persistence. As shown in Fig. 3e, core segments exhibit the highest orientational persistence, consistent with extended, radially biased conformations. Furthermore, this persistence increases with condensate age, correlating with the increase in *R*_*g*_. In contrast, segments in the outer shell show weak orientational correlations, reflecting frequent directional changes and more flexible, partially folded conformations. Together, these results establish a direct connection between spatially heterogeneous chain conformations and dynamics: extended, aligned RNA chains in the condensate core experience stronger topological constraints and slower rearrangements, while the more compact and flexible chains at the periphery enable faster, liquid-like dynamics.

### D. Molecular Origins of Condensate Aging

Due to the expansion and preferential orientation of RNA chains near the condensate core, we posited that chain segments in this region approach one another to form dense, mesh-like networks,^84^ thereby impeding relaxation, which would result in accelerating aging. To quantitatively characterize the network architecture, we employ primitive path analysis (PPA), used in polymer physics to probe the topology of entanglement networks.^83,85^ In PPA, each polymer chain, with fixed endpoints, is contracted to its shortest contour length without crossing the neighboring chains. This yields a primitive path composed of straight segments connected by kinks.^85–87^ Each kink represents a topological constraint imposed by the surrounding chains. The number of kinks per chain, *Z*, provides a measure of entanglement density. The entanglement length is given by *N*_*e*_ ≈ *N/Z* for a polymer of length *N* . Regions with higher kink densities experience stronger topological confinement, leading to restricted segmental motion and longer relaxation times.^88,89^

We find that the average number of kinks per chain increases steadily with condensate age, growing from *Z* ∼ 5.0 at early times to *Z* ∼ 8.0 at later stages (Fig. 4a), indicating the progressive buildup of a topologically constrained network. Perhaps not surprisingly, this is highly correlated to the increase of chain density in the condensate, except during the initial phase of growth. It is clear that when *Z* and condensate density track each other at long times, there is a marked change in the network architecture (Fig. 4a). Remarkably, the kinks are strongly enriched in the core, where their density is approximately 5 times higher than in the outer shell (Fig. 4b). (We quantify this spatial variation using the local kink density *ρ*_*z*_ = *Z*_shell_*/V*_shell_, where *Z*_shell_ is the number of kinks and *V*_shell_ is the volume of a given radial shell.) As a result, the local plateau modulus 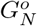, estimated using 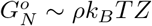 with *ρ* denoting the nucleotide density, is maximal in the core region (Fig. 4c). These results demonstrate that RNA chains near the core are significantly more entangled, likely due to their extended conformations and the higher local nucleotide density (Fig. S21). The resulting increase in network crosslinking (Fig. S22) confers greater mechanical rigidity to the core, strongly suppressing relaxation and accelerating aging relative to the surface.

**FIG. 4.**
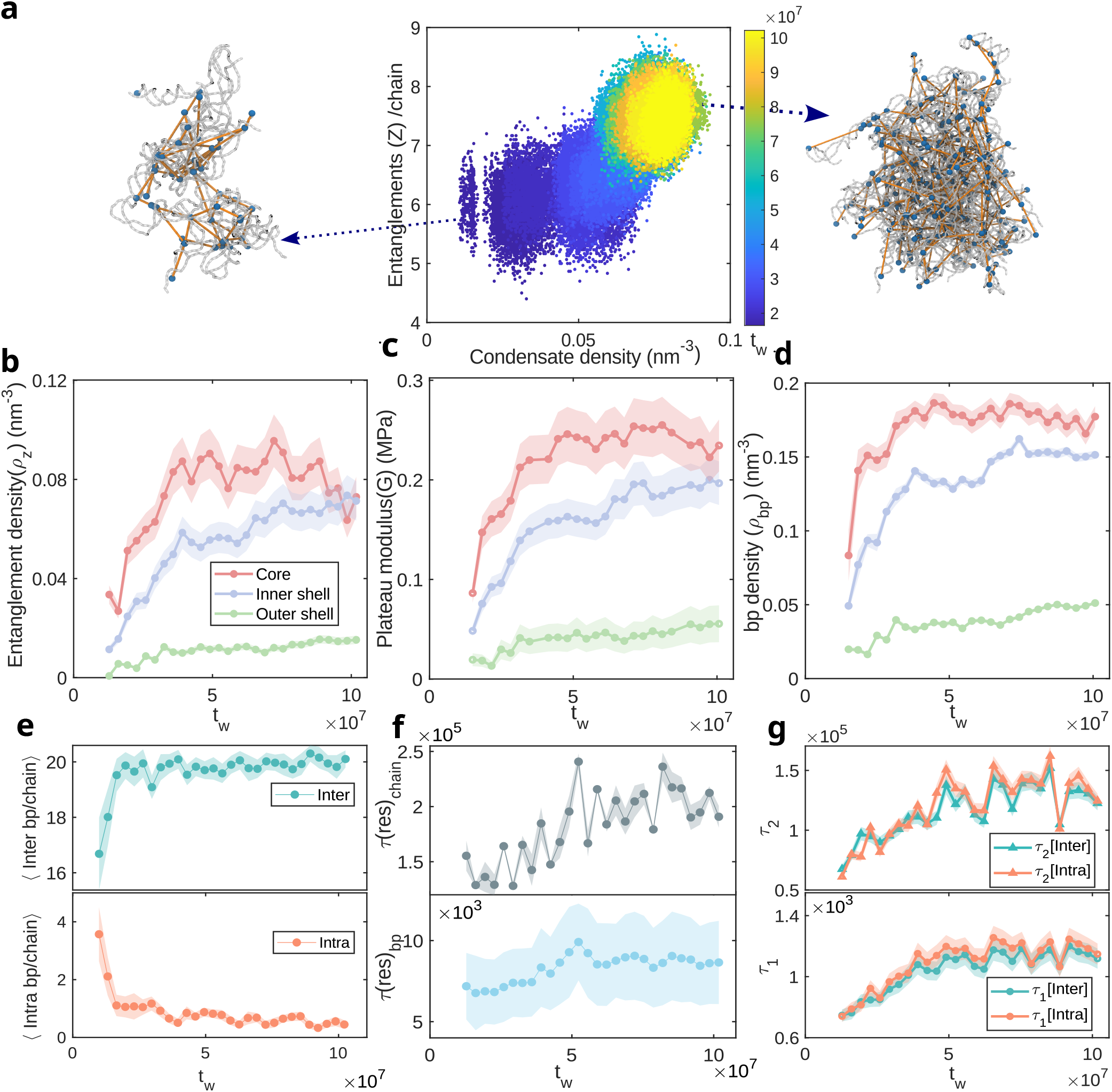
Molecular basis of spatial heterogeneity in aging of CAG-repeat RNA condensates. (a) Evolution of the condensate internal density and entanglement as a function of time. Representative snapshots from primitive path analysis illustrating the evolution of the topological network structure as the droplet ages. Blue beads indicate entanglement points (kinks) representing topological constraints imposed by the neighboring chains. Yellow lines show primitive path segments connecting successive kinks. The condensate transitions from a sparse network (left) to a densely entangled network (right) upon aging. (b) Spatial and temporal evolution of entanglement (kink) density across radial shells demonstrates that the core develops a denser mesh-like network as the condensate ages compared to outer shells. (c) Plateau modulus 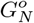 showing the core region exhibits the highest values, indicating greater network crosslinking and higher mechanical rigidity compared to the periphery. (d) Watson–Crick base-pair density shows a sharp core-to-periphery gradient with approximately ∼ 4-fold enrichment from the outer shell to the core. (e) Decomposition of base-pairing into intra-(orange) and inter-chain (green) contributions reveals that inter-chain base-pairing dominates and primarily drives the network formation and aging dynamics. (f) Both base-pair residence time and chain-chain contact lifetime increase progressively as the condensate ages, indicating strengthening of the network through longer-lived molecular interactions. (g) Segmental relaxation timescales partitioned into intra- and inter-chain contributions show comparable local dynamics.

To identify the molecular interactions that stabilize the dense topological network, we next analyze the distribution of canonical Watson–Crick base pairs (bp). We calculate the bp density in each shell as *ρ*(bp)_shell_ = *N* (bp)_shell_*/V*_shell_, where *N* (bp)_shell_ is the number of bps formed within a given shell at age *t*_*w*_. The bp density exhibits a similar gradient to the kink density, peaking in the core and spanning nearly a 4-fold variation across the droplet radius (Fig. 4d). Most importantly, *ρ*_bp_ increases monotonically with condensate age before saturating at long times, indicating continual densification of the bp interaction network during aging. Together with the growing entanglement network, this pronounced bp density gradient provides a direct molecular explanation for the emergence of core-shell organization. Extensive bp in the core facilitates chain unfolding and expansion, which in turn increases inter-chain entanglement and further constrains molecular motion. This leads to progressively longer relaxation times and accelerated aging in the core. Conversely, the periphery is characterized by a sparse and transient bp network, allowing rapid rearrangements and preserving liquid-like dynamics.

To further dissect the nature of the bp interactions, we decompose them into intra- and inter-chain contributions and track their evolution with condensate age. Intra-chain bp remains minimal (0-1 bps per chain) and decreases over time (Fig. 4e), consistent with the energetic and entropic penalties associated with forming secondary structures such as hairpins in a densely crowded environment.^49,90,91^ In contrast, inter-chain bp dominates the interaction network, with each chain participating in approximately 19-20 bps. This dominance reflects the thermodynamic advantage of intermolecular pairing between extended RNA chains^49^, which can form complementary contacts with minimal conformational cost.

Finally, we find that aging is accompanied not only by an increase in the number of bps but also by enhanced persistence of these interactions. To quantify this effect, we calculate two lifetimes: i) the bp resident time, *τ* (res)_bp_, defined as the mean duration over which an individual bp remains intact, and ii) the chain-chain contact lifetime, *τ* (res)_chain_, defined as the total time during which two RNA molecules remain connected by at least one bp. Both lifetimes increase modestly with condensate age (Fig. 4f), demonstrating that the homotypic RNA interaction network becomes increasingly more stable and long-lived at both the bp and whole-chain level.

### E. Aging Depends on RNA Sequence

The emergence of a robust, spatially organized core-shell network during condensate aging is fundamentally driven by sequence-specific G-C bp interactions for the CAG repeat. Recent studies have shown that sequence periodicity plays a critical role in governing RNA phase behavior: scrambled sequences with identical nucleotide composition but lacking repeat complementarity display a significantly reduced propensity for phase separation.^49,57,92–94^ To elucidate how RNA sequence modulates aging dynamics and separate the contribution of density and entanglement, we directly compare the condensate evolution of CAG-repeat RNA with that of a scrambled sequence (Figs. 5a-d).

**FIG. 5.**
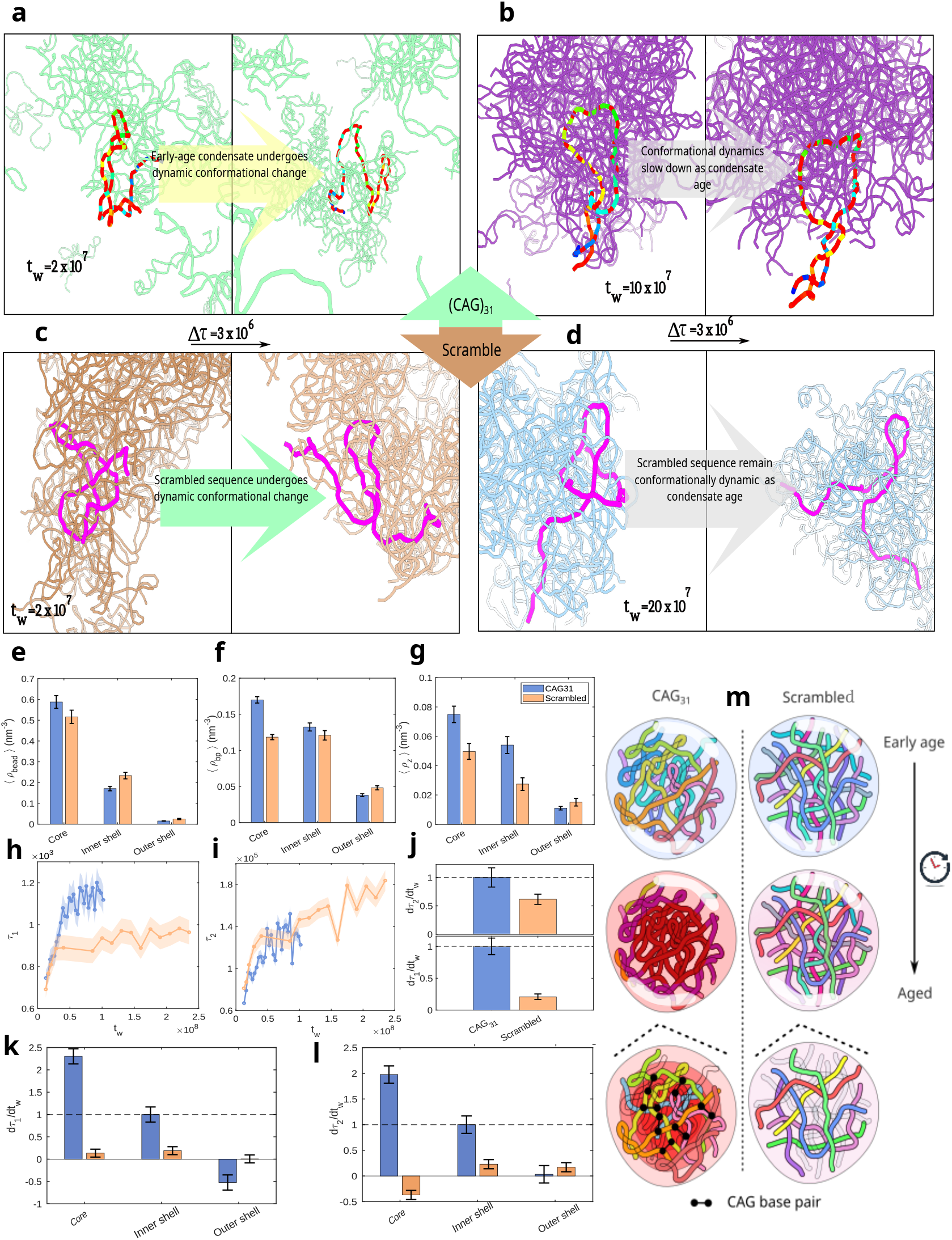
Sequence effects on condensate aging: (a) At early age (*t*_*w*_ = 2 *×* 10^7^ timesteps), the highlighted (CAG)_31_ chain undergoes large-amplitude conformational changes over Δ*τ* = 3 *×* 10^6^ timesteps, indicating absence of structural memory. (b) At mature age (*tw* = 10 *×* 10^7^ timesteps), the same chain retains its conformation across Δ*τ*, signaling the emergence of structural memory driven by persistent inter-chain base pairing. (c) The scrambled chain is equally dynamic at early age (*t*_*w*_ = 2 *×* 10^7^ timesteps). (d) Unlike (CAG)_31_, the scrambled chain remains conformationally mobile even at mature age (*tw* = 20 *×* 10^7^ timesteps), confirming the absence of structural memory. (e) Overall nucleotide concentrations ⟨*ρ*_bead_⟩ across shells show comparable spatial patterns for both sequences. (f) Base-pair density comparison between the two droplets: CAG-repeat displays pronounced base-pair density gradient from core to outer shell, while the scrambled droplet shows flat distribution in bp density between core and inner shell. (g) Entanglement kink density for CAG-repeat and scrambled sequences. The scrambled condensate exhibits systematically lower kink density from core to outer shell. (h) The fast relaxation time (*τ*_1_) of the scrambled condensate shows modest initial rise followed by saturation at lower values compared to those of the CAG-repeat. (i) Same as (h) but for the slow relaxation time (*τ*_2_). (j) The relative rates of change for the two relaxation times show the CAG-repeat condensate ages faster than the scrambled system. (k) and (l), Shell-wise relative aging rate *d*⟨*τ*_1_⟩ */dt*_*w*_ and *d*⟨*τ*_2_⟩ */dt*_*w*_ reveals that the CAG-repeat condensate exhibit clear spatial aging while the scramble sequence ages much slower and more homogeneously. (m) The schematic illustrates the underlying molecular contrast: with age, (CAG)_31_ develops a compact entangled core stabilized by persistent C–G inter-chain base pairs, while the scrambled condensate remains loosely organized despite identical nucleotide composition.

Topological entanglement analysis reveals striking sequence-dependent differences in spatial organization of the two condensates (Fig. 5g). While the CAG-repeat condensate exhibits the core-shell gradient in the kink density described above, the scrambled sequence shows a substantially lower number of kinks in both the core and inner shell. This reduction indicates that the scrambled sequence does not form a dense, spatially stratified network of topological constraints comparable to that of the CAG-repeat condensate. Importantly, this difference cannot be attributed solely to variations in molecular density, as both the condensates exhibit similar values, with their core equally densely packed (Fig. 5e). Despite this, the scrambled condensate does not form a strongly entangled network, demonstrating that high local concentration alone is insufficient to drive condensate aging in the absence of sequence complementarity.

Direct comparison of bp density provides the clearest evidence for sequence-specific driving forces underlying network formation (Fig. 5f). The CAG-repeat condensate exhibits a pronounced bp density gradient, with maximal bp in the core that decays monotonically toward the periphery. In contrast, the scrambled condensate exhibits nearly uniform bp density between the core and inner shell, with little spatial heterogeneity. The absence of a strong bp gradient explains both the lack of a core-shell architecture and the reduced topological entanglement in the scrambled condensate. These results show that the scrambled sequence distributes bp interactions more uniformly and non-cooperatively, in contrast to the highly cooperative network formed by the CAG repeat. As a result, the scrambled sequence requires higher critical concentrations to undergo phase separation, in agreement with experimental observations.^13^

Consistent with the absence of a stable, sequence-specific bp network, the scrambled condensate exhibits significantly slower aging dynamics than the CAG-repeat system. While the CAG-repeat droplet shows rapid early aging marked by a sharp increase in relaxation times, the scrambled condensate exhibits only a modest initial increase followed by an early plateau in the fast relaxation time *τ*_1_ at significantly lower value (Fig. 5h). This behavior shows that the local cages restricting short-time segmental motion in the scrambled network do not progressively strengthen with age, in contrast to the CAG repeat. Interestingly, the slow relaxation time *τ*_2_ evolves similarly in both systems, suggesting that this mode may be less sensitive to network architecture and instead reflect more global droplet-scale rearrangements (Fig. 5i). Nevertheless, the overall aging rate of the scramble condensate is ∼ 4-5 times smaller than that of the CAG-repeat droplet (Fig. 5j). As a result, the scrambled condensate exhibits minimal spatial differentiation in aging behavior, with nearly uniform aging across radial shells (Figs. 5k and l).

We also perform simulations of a longer repeat sequence, (CAG)_47_, to examine how RNA length influences condensate aging relative to (CAG)_31_. Despite comparable nucleotide densities within the condensates (Fig. S29), condensates formed by longer (CAG)_47_ chains exhibit longer relaxation times and age more rapidly due to their greater propensity for topological entanglement (Figs. S30, S31, and S32). Taken together, our results demonstrate that sequence-specific base pair formation is essential for the emergence of spatially heterogeneous, progressively aging RNA condensates. In the absence of repeat complementarity, RNA networks are weakly connected, spatially uniform, and largely resistant to long-term dynamical arrest.

## III. DISCUSSIONS AND CONCLUSION

In this work, we demonstrate that aging of RNA condensates emerge naturally from sequence-encoded interactions and polymer connectivity, without requiring *ad hoc* tuning of interaction parameters. Our simulations reveal that aging in RNA condensate is not spatially uniform, but instead proceeds heterogeneously, giving rise to a progressively arrested, solid-like core surrounded by a relatively fluid-like shell. The emergent spatiotemporal heterogeneity, and the resulting dependence of the dynamics on the distance from the droplet center provides a unifying framework that links molecular-scale RNA interactions to mesoscale material transitions in pathological RNA condensates.

Another key finding is that condensate aging is strongly sequence dependent. CAG-repeat RNAs, which engage in extensive homotypic base pairing, undergo pronounced aging characterized by a progressive slowing of dynamics, the emergence of long-lived structural memory, and the buildup of a dense interaction network. In contrast, scrambled RNAs with identical nucleotide composition but lacking repeat complementarity show minimal aging, weak spatial heterogeneity, and largely preserved liquid-like behavior. These results reinforce the notion that RNA sequence pattern and structure, not merely composition, play a critical role in dictating the condensate material properties,^8,95^ extending previous observations on phase separation propensity to condensate aging.

The emergence of multiple relaxation timescales and their evolution with condensate age places RNA condensates in close analogy with aging soft materials, such as polymer gels, random copolymers, colloidal glasses, and jammed networks.^72,77,78,96–100^ In this context, the overlap function and subdiffusive motion reveal that RNA condensates gradually lose ergodicity, retaining memory of initial configurations over increasingly long timescales. Importantly, our shell-resolved analyses show that the loss of ergodicity is spatially localized: aging is most rapid in the condensate core and slowest near the interface. This observation suggests that condensate aging should be viewed as a spatially heterogeneous process rather than a uniform bulk transition.

Could aging be a generic feature of naturally evolved RNA sequences? Although it is difficult to answer this question in complete generality, our work suggests a possibility based on two criteria that must be satisfied for aging. First, the propensity to form condensates is encoded in the monomer free energy spectra.^101,102^ In the repeat RNA sequences, condensates form readily because the free energy gap between the ground and excited states is small (on the order of *k*_*B*_*T*). Second, upon entering the droplet, the RNA chains have to unfold to facilitate multiple intermolecular interactions, leading to entanglement, which over time would result in aging. However, if the RNA is highly structured, unfolding in the condensate would not occur readily,^103^ which is required for extensive intermolecular interactions. Such sequences (mRNAs for example), while capable of forming condensates, are unlikely to display age-dependent dynamics.

Our results provide a mechanism for the spontaneous emergence of core-shell architecture during RNA condensate aging: RNAs near the core unwind and undergo a transition from hairpin to extended conformations, align preferentially along the radial direction, and form dense networks of inter-chain base pairs. These structural changes result in topological entanglement, leading to a mechanically stiff, slowly relaxing core.^83,104^ In contrast, RNAs near the periphery remain more compact, weakly entangled, and dynamically flexible, preserving liquid-like behavior. This architecture arises naturally from the coupling between RNA conformations, base-pairing interactions, and chain topology.^105^ Importantly, the core-shell organization does not require explicit energetic heterogeneity or multiphase coexistence. Instead, it emerges naturally from self-reinforcing feedback: base pairing drives chain expansion; expansion increases entanglement; entanglement slows relaxation; and slow relaxation reinforces network growth. Such a feedback loop maybe a general physical principle underlying condensate aging across diverse biological systems.

The core-shell architecture uncovered here, with solid-like interior enclosed by a fluid and dynamic shell, has been observed in previous computational studies in a different context.^10,35,36,56^ In particular, CG simulations of multi-domain FUS condensates^35^ showed the formation of an orientationally ordered, fibril-like arrested core surrounded by a disordered liquid-like shell. The findings in FUS^35^ and results presented here suggest that the aging mechanisms in biopolymers with vastly different sequences and interactions could be similar. Indeed, such a structure is also in qualitative agreement with the behavior of a telomeric repeat RNA^27^ which undergoes a liquid-to-solid transition in the condensate core. A similar architecture has also been suggested for the nucleolus which features an extensively entangled core and a more fluid shell.^106^ Stress granules, which are predominantly RNA–protein networks, form multiple stable cores surrounded by a dynamic shell.^32,107^

Despite the common features noted above, it is important to stress that there are other kinds of architecture.^74^ NMR and Raman spectroscopic studies of FUS condensates have revealed that aging drives the nucleation of a solid fibrillar shell surrounding a presumably more liquid-like core.^25,108–110^ Similar observations have been made for hnRNPA1, where fibril formation is specifically promoted at the interface of the two phases.^111–113^ In cells, P granules harbor a dynamic liquid core enclosed within a gel-like, stable shell,^114^ while nuclear speckles adopt a complex multi-layered architecture.^115^Our simulations suggest a mechanistic bridge between these architectures and molecular-scale interactions, indicating that spatial heterogeneity in condensates can arise from intrinsic network formation in addition to external regulators or compositional gradients.^116^

The aging behavior observed here has direct implications for RNA biology,^84,117^ particularly in the context of repeat expansion disorders. Expanded CAG-repeat RNAs are implicated in several neurodegenerative diseases, where they form persistent nuclear foci and sequester RNA-binding proteins. Our results suggest that sequence-driven condensate aging may contribute to the pathological persistence of such RNA assemblies by favoring irreversible network formation and dynamic arrest. The gradual solidification of the core may reduce molecular exchange and impair normal RNA processing. More broadly, our findings highlight how RNA sequence can encode not only condensate formation but also long-term material evolution.^118,119^ This raises the possibility that cells may exploit or regulate condensate aging through sequence design, RNA modifications, helicase activity, or changes in ionic conditions and temperature to maintain functional condensates in a dynamic and reversible state. Conversely, failure to regulate these processes could drive aberrant aging and pathological aggregation.

Our results also highlight limitations inherent to computational approaches. While CG simulations enable access to mesoscale structure and long-time relaxation processes, they remain constrained by finite system sizes, timescales and details of microscopic interactions. Nonetheless, the emergence of aging, spatial heterogeneity, and sequence dependence suggests that these phenomena are robust and likely persist beyond the simple model employed here. Future multiscale approaches combining CG modeling with experimental measurements will be essential to fully characterize condensate aging across biologically relevant timescales. From a physics perspective, this work emphasizes the importance of treating condensates as evolving polymer networks rather than equilibrium liquid droplets. Aging, spatial heterogeneity, and nonergodicity emerge naturally when interaction networks grow and reorganize over time. These features challenge simplified descriptions of condensates based solely on surface tension or bulk viscosity and call for theoretical frameworks that integrate polymer physics, network theory, and nonequilibrium dynamics.

## IV. METHODS

Following the Single-Interaction-Site (SIS) model introduced by Nguyen *et al*.^49^, each nucleotide is mapped to a single bead carrying the base identity (A, G or C). Albeit simple, the SIS model preserves sequence specificity while permitting multi-chain simulations over the timescales required to investigate condensate aging. The total energy of an *N* –chain RNA system is

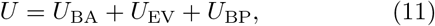

where the three contributions are described below.

### Backbone Connectivity (*U*_BA_)

Backbone integrity is enforced by harmonic restraints between successive bond lengths and angles:

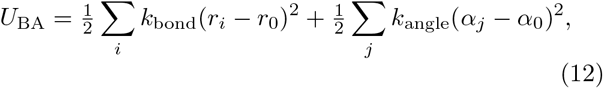

where *k*_bond_ = 15.0 kcal mol^−1 Å−2^, *r*_0_ = 5.9 ^Å^, *k*_angle_ = 10.0 kcal mol^−1^ rad^−2^, *α*_0_ = 2.618 rad.

### Excluded Volume (*U*_EV_)

Short–range steric repulsion is treated using the Weeks–Chandler–Andersen potential:

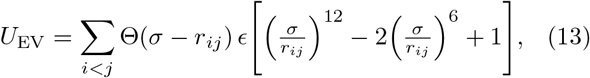

where *σ* = 10.0 ^Å^ and *ϵ* = 2.0 kcal mol^−1^. As in the original model, this term is only calculated for pairs that are not involved in bond or angle restraints, and are separated by at least two other beads along the chain.

### Base–Pairing (*U*_BP_)

Canonical Watson–Crick pairing between complementary nucleotides is represented by the many–body potentia:

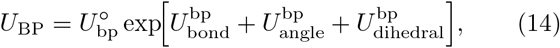

where

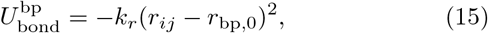

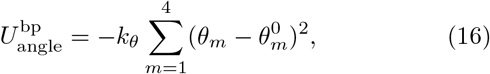

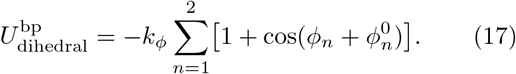

The parameters in the equation above are *r*_bp,0_ = 13.8 ^Å^, *k*_*r*_ = 3.0 ^Å−2^, *k*_*θ*_ = 1.5 rad^−2^, *k*_*ϕ*_ = 0.5, with angular references 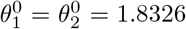 rad and 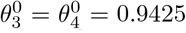 rad, and dihedral references 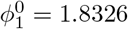 rad, 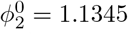 rad. The energy scale 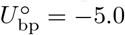 kcal mol^−1^. Additional details of the SIS model maybe found elsewhere^49^.

### Simulation Protocol

We place 64 chains of (CAG)_31_ in a cubic periodic box whose edge was chosen to match the desired bulk concentration (20–500 µM). Langevin dynamics with a reduced friction of *γ* = 0.01 ps^−1^ was propagated using a 10 fs time step in OpenMM 7.7. The thermostat is maintained at *T* = 300 K. Each production run lasted 2 *×* 10^8^ steps (≈2 ms in reduced time units).

The scrambled sequence is the one used previously:^49^

GCACGGGCACACGCCGGACAGACCAGAGACGGCCAAGGACGCAGCAGAGCACCGACGAGCGCAGCCGAGAAGCGCGCAGCCGGCGACGGCAGC

### Cluster Identification

A single-linkage algorithm groups chains whose minimum inter-nucleotide distance is 16 ^Å^. This threshold is ≈1.3 *r*_bp,0_, accounting for the fluctuation in base pairing distance.

### Criterion for Base-Pair formation

A base pair is deemed present when its interaction energy satisfies

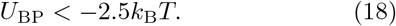

This purely energetic definition automatically enforces the distance and orientational requirements embodied in *U*_BP_.

### Droplet Selection for Aging Analysis

We perform simulations at various RNA concentrations: 400, 200 and 100 *µ*M. Only well-formed condensates are selected for aging analyses. In particular, for (CAG)_31_ at 400 *µ*M, we select the first frame in which the largest cluster contained *at least* 30 chains. For the scrambled sequence, we start the analysis when the droplet size reaches *at least* 35 chains. Thereafter, all analyses focus exclusively on the interior of that growing condensate. As the condensate accrues additional chains it eventually reaches the simulation-limited maximum of 64 chains for both systems. We run much longer simulations for the scrambled droplet compared to (CAG)_31_ (a total of 2.54 × 10^8^ timesteps) to examine whether aging continues at long times. In more dilute conditions, condensate formation is either incomplete (smaller droplets with fewer chains at 200 *µ*M) or absent (100 *µ*M). This is presumably due to the diffusion-limited nature of condensate growth. Thus we did not choose those for the aging analyses.

The interval is partitioned into *n* = 27 contiguous windows. Each window spans ≈3.3 × 10^6^ time steps (≈0.033 ms), yielding uniform statistics across the aging trajectory.

### Aging rate estimation

To quantify the rate of change (slope) in the relaxation behavior (*τ*_1_, *τ*_2_), and size of RNA chains (*R*_*g*_, *ν*), we fit to a linear model:

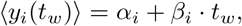

where *y*_*i*_(*t*_*w*_) is the observable of interest at age *t*_*w*_ for each region *i* (core, inner shell, outer shell) independently. Here, *α*_*i*_ is the intercept (initial value of y-axis) and *β*_*i*_ is the slope (aging rate with units timestep^−1^). Parameters were estimated using standard least-squares minimization.

To rigorously quantify the uncertainty in estimated aging rates and assess their statistical significance, we computed 95% confidence intervals for each linear regression slope using the maximum likelihood framework. Specifically, for each fit we employed the Jacobian matrix of residuals to calculate the covariance matrix of the fitted parameters using:

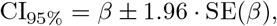

where SE(*β*) is the standard error of the slope estimate derived from the covariance matrix diagonal. This approach is implemented via MATLAB’s nlnparci function.

The quality of the linear fit was further verified by calculating the coefficient of determination *R*^2^ for each regression. All fits yielded *R*^2^ *>* 0.90, confirming that linear model adequately describes the temporal evolution of the observable of interest. No smoothing, filtering, or data binning was applied prior to regression analysis.

## V. ACKNOWLEDGEMENTS

We gratefully acknowledge support from the University at Buffalo to H.T.N. Computational support is provided by the Center for Computational Research at the University at Buffalo. D.T. is grateful to the National Science Foundation (CHE 2320256) and the Welch Foundation through the Collie-Welch Chair (F-0019) for support.

## Notes

### Competing Interest Statement

The authors have declared no competing interest.

## References

1 C. P. Brangwynne, C. R. Eckmann, D. S. Courson, A. Rybarska, C. Hoege, J. Gharakhani, F. Jülicher, and A. A. Hyman, Science 324, 1729 (2009), https://www.science.org/doi/pdf/10.1126/science.1172046.

2 S. F. Banani, H. O. Lee, A. A. Hyman, and M. K. Rosen, Nature Reviews Molecular Cell Biology 18, 285–298 (2017).

3 A. A. Hyman, C. A. Weber, and F. Jülicher, Annual Review of Cell and Developmental Biology 30, 39–58 (2014).

4 S. Alberti and A. A. Hyman, Nature Reviews Molecular Cell Biology 22, 196–213 (2021).

5 A. Amir, Y. Oreg, and Y. Imry, Proceedings of the National Academy of Sciences 109, 1850 (2012), https://www.pnas.org/doi/pdf/10.1073/pnas.1120147109.

6 L. Jawerth, E. Fischer-Friedrich, S. Saha, J. Wang, T. Franzmann, X. Zhang, J. Sachweh, M. Ruer, M. Ijavi, S. Saha, J. Mahamid, A. A. Hyman, and F. Jülicher, Science 370, 1317 (2020), https://www.science.org/doi/pdf/10.1126/science.aaw4951.

7 R. Takaki, L. Jawerth, M. Popović, and F. Jülicher, PRX Life 1, 013006 (2023).

8 I. Alshareedah, W. M. Borcherds, S. R. Cohen, A. Singh, A. E. Posey, M. Farag, A. Bremer, G. W. Strout, D. T. Tomares, R. V. Pappu, T. Mittag, and P. R. Banerjee, Nature Physics 20, 1482–1491 (2024).

9 Patel, H. Lee, L. Jawerth, S. Maharana, M. Jahnel, M. Hein, S. Stoynov, J. Mahamid, S. Saha, T. Franzmann, A. Pozniakovski, I. Poser, N. Maghelli, L. Royer, M. Weigert, E. Myers, S. Grill, D. Drechsel, A. Hyman, and S. Alberti, Cell 162, 1066 (2015).

10 A. Garaizar, J. R. Espinosa, J. A. Joseph, G. Krainer, Y. Shen, T. P. Knowles, and R. Collepardo-Guevara, Proceedings of the National Academy of Sciences 119 (2022), 10.1073/pnas.2119800119.

11 L. Berthier and G. Biroli, Rev. Mod. Phys. 83, 587 (2011).

12 T. R. Kirkpatrick and D. Thirumalai, Rev. Mod. Phys. 87, 183 (2015).

13 A. Jain and R. D. Vale, Nature 546, 243 (2017).

14 M. M. Fay, P. J. Anderson, and P. Ivanov, Cell Reports 21, 3573–3584 (2017).

15 C. Mathieu, R. V. Pappu, and J. P. Taylor, Science 370, 56 (2020), https://www.science.org/doi/pdf/10.1126/science.abb8032.

16 M. Linsenmeier, M. Hondele, F. Grigolato, E. Secchi, K. Weis, and P. Arosio, Nature Communications 13 (2022), 10.1038/s41467-022-30521-2.

17 Y. Pan, J. Lu, X. Feng, S. Lu, Y. Yang, G. Yang, S. Tan, L. Wang, P. Li, S. Luo, and B. Lu, Nature Chemical Biology 19, 1372–1383 (2023).

18 S. Wegmann, B. Eftekharzadeh, K. Tepper, K. M. Zoltowska, R. E. Bennett, S. Dujardin, P. R. Laskowski, D. MacKenzie, T. Kamath, C. Commins, C. Vanderburg, A. D. Roe, Z. Fan, A. M. Molliex, A. Hernandez-Vega, D. Muller, A. A. Hyman, E. Mandelkow, J. P. Taylor, and B. T. Hyman, The EMBO Journal 37 (2018), 10.15252/embj.201798049.

19 S. Jeon, Y. Jeon, J.-Y. Lim, Y. Kim, B. Cha, and W. Kim, Signal Transduction and Targeted Therapy 10 (2025), 10.1038/s41392-024-02070-1.

20 H. E. Castillo, C. Chamon, L. F. Cugliandolo, and M. P. Kennett, Phys. Rev. Lett. 88, 237201 (2002).

21 Y. Lin, D. Protter, M. Rosen, and R. Parker, Molecular Cell 60, 208 (2015).

22 N. A. Erkamp, T. Sneideris, H. Ausserwöger, D. Qian, S. Qamar, J. Nixon-Abell, P. St George-Hyslop, J. D. Schmit, D. A. Weitz, and T. P. J. Knowles, Nature Communications 14 (2023), 10.1038/s41467-023-36059-1.

23 S. K. Rai, R. Khanna, A. Avni, and S. Mukhopadhyay, Proceedings of the National Academy of Sciences 120, e2216338120 (2023), https://www.pnas.org/doi/pdf/10.1073/pnas.2216338120.

24 K. Makasewicz, T. N. Schneider, P. Mathur, S. Stavrakis, A. J. deMello, and P. Arosio, Biophysical Journal 124, 115–124 (2025).

25 Y. Shen, A. Chen, W. Wang, Y. Shen, F. S. Ruggeri, S. Aime, Z. Wang, S. Qamar, J. R. Espinosa, A. Garaizar, P. S. George-Hyslop, R. Collepardo-Guevara, D. A. Weitz, D. Vigolo, and T. P. J. Knowles, Proceedings of the National Academy of Sciences 120, e2301366120 (2023), https://www.pnas.org/doi/pdf/10.1073/pnas.2301366120.

26 W. Yu, X. Guo, Y. Xia, Y. Ma, Z. Tong, L. Yang, X. Song, R. N. Zare, G. Hong, and Y. Dai, Nature Chemistry 17, 756–766 (2025).

27 T. S. Mahendran, G. M. Wadsworth, A. Singh, R. Gupta, and P. R. Banerjee, Nat. Chem., 1236 (2025).

28 M. Feric, N. Vaidya, T. S. Harmon, D. M. Mitrea, L. Zhu, T. M. Richardson, R. W. Kriwacki, R. V. Pappu, and C. P. Brangwynne, Cell 165, 1686 (2016).

29 S. Choi, M. O. Meyer, P. C. Bevilacqua, and C. D. Keating, Nature Chemistry 14, 1110–1117 (2022).

30 D. S. Protter and R. Parker, Trends in Cell Biology 26, 668 (2016).

31 A. Z. Lin, K. M. Ruff, F. Dar, A. Jalihal, M. R. King, J. M. Lalmansingh, A. E. Posey, N. A. Erkamp, I. Seim, A. S. Gladfelter, and R. V. Pappu, Nature Communications 14 (2023), 10.1038/s41467-023-43489-4.

32 D. W. Sanders, N. Kedersha, D. S. Lee, A. R. Strom, V. Drake, J. A. Riback, D. Bracha, J. M. Eeftens, A. Iwanicki, A. Wang, M.-T. Wei, G. Whitney, S. M. Lyons, P. Anderson, W. M. Jacobs, P. Ivanov, and C. P. Brangwynne, Cell 181, 306 (2020).

33 S. Saurabh, T. N. Chong, C. Bayas, Dahlberg, H. N. Cartwright, W. E. Moerner, . and L. P. Shapiro, Science Advances 8, eabm6570 (2022), https://www.science.org/doi/pdf/10.1126/sciadv.abm6570.

34 A. S. Holehouse and S. Alberti, Molecular Cell 85, 290–308 (2025).

35 S. Ranganathan and E. Shakhnovich, Biophysical Journal 121, 2751 (2022).

36 S. Biswas and D. A. Potoyan, PRX Life 2, 023011 (2024).

37 C. Moore, E. Wong, U. Kaur, U. S. Chio, Z. Zhou, M. Ostrowski, K. Wu, I. Irkliyenko, S. Wang, V. Ramani, and G. J. Narlikar, Science 390, eadr0018 (2025), https://www.science.org/doi/pdf/10.1126/science.adr0018.

38 W. Kang, Z. Wu, X. Huang, H. Qi, J. Wu, J. Wang, J. Li, S. Wu, B.-H. Kang, B. Li, J. Ma, and C. Xue, Nature Communications 16 (2025), 10.1038/s41467-025-62074-5.

39 S. Ranganathan, P. Dasmeh, S. Furniss, and E. Shakhnovich, Proceedings of the National Academy of Sciences 120, e2215828120 (2023), https://www.pnas.org/doi/pdf/10.1073/pnas.2215828120.

40 A. Patel, L. Malinovska, S. Saha, J. Wang, S. Alberti, Y. Krishnan, and A. A. Hyman, Science 356, 753 (2017), https://www.science.org/doi/pdf/10.1126/science.aaf6846.

41 A. Ghosh, D. Kota, and H.-X. Zhou, Nature Communications 12 (2021), 10.1038/s41467-021-26274-z.

42 D. Zwicker, R. Seyboldt, C. A. Weber, A. A. Hyman, and F. Jülicher, Nature Physics 13, 408–413 (2016).

43 T. S. Harmon, A. S. Holehouse, M. K. Rosen, and R. V. Pappu, eLife 6, e30294 (2017).

44 G. L. Dignon, W. Zheng, R. B. Best, Y. C. Kim, and J. Mittal, Proceedings of the National Academy of Sciences 115, 9929 (2018), https://www.pnas.org/doi/pdf/10.1073/pnas.1804177115.

45 J. McCarty, K. T. Delaney, S. P. O. Danielsen, G. H. Fredrickson, and J.-E. Shea, The Journal of Physical Chemistry Letters 10, 1644 (2019), 10.1021/acs.jpclett.9b00099.

46 S. Ranganathan and E. I. Shakhnovich, eLife 9, e56159 (2020).

47 G. Tesei, T. K. Schulze, R. Crehuet, and K. Lindorff-Larsen, Proceedings of the National Academy of Sciences 118, e2111696118 (2021), https://www.pnas.org/doi/pdf/10.1073/pnas.2111696118.

48 J. A. Joseph, A. Reinhardt, A. Aguirre, P. Y. Chew, K. O. Russell, J. R. Espinosa, A. Garaizar, and R. Collepardo-Guevara, Nature Computational Science 1, 732–743 (2021).

49 H. T. Nguyen, N. Hori, and D. Thirumalai, Nat. Chem. 14, 775 (2022).

50 K. L. Saar, D. Qian, L. L. Good, A. S. Morgunov, R. Collepardo-Guevara, R. B. Best, and T. P. J. Knowles, Chemical Reviews 123, 8988 (2023), pMID: 37171907, 10.1021/acs.chemrev.2c00586.

51 B. Szala-Mendyk, T. M. Phan, P. Mohanty, and J. Mittal, Current Opinion in Chemical Biology 75, 102333 (2023).

52 W. Zheng, G. L. Dignon, N. Jovic, X. Xu, R. M. Regy, N. L. Fawzi, Y. C. Kim, R. B. Best, and J. Mittal, The Journal of Physical Chemistry B 124, 11671 (2020), pMID: 33302617, 10.1021/acs.jpcb.0c10489.

53 K. A. Lorenz-Ochoa and C. R. Baiz, Journal of the American Chemical Society 145, 27800–27809 (2023).

54 A. R. Tejedor, R. Collepardo-Guevara, J. Ramírez, and J. R. Espinosa, The Journal of Physical Chemistry B 127, 4441–4459 (2023).

55 D. Sundaravadivelu Devarajan, J. Wang, B. Szala-Mendyk, S. Rekhi, A. Nikoubashman, Y. C. Kim, and J. Mittal, Nature Communications 15 (2024), 10.1038/s41467-024-46223-w.

56 A. R. Tejedor, I. Sanchez-Burgos, M. Estevez-Espinosa, A. Garaizar, R. Collepardo-Guevara, J. Ramirez, and J. R. Espinosa, Nature Communications 13 (2022), 10.1038/s41467-022-32874-0.

57 R. Takaki and D. Thirumalai, Proceedings of the National Academy of Sciences 121, e2409973121 (2024), https://www.pnas.org/doi/pdf/10.1073/pnas.2409973121.

58 H. T. Orr and H. Y. Zoghbi, Annual Review of Neuroscience 30, 575 (2007).

59 J. Schmoll, M. Novakovic, and F. H.-T. Allain, Nature Chemistry 17, 1785–1794 (2025).

60 Z. Guo and D. Thirumalai, Biopolymers 36, 83 (1995).

61 A. S. Keys, L. O. Hedges, J. P. Garrahan, S. C. Glotzer, and D. Chandler, Phys. Rev. X 1, 021013 (2011).

62 S. S. Schoenholz, E. D. Cubuk, D. M. Sussman, E. Kaxiras, and A. J. Liu, Nature Physics 12, 469–471 (2016).

63 G. B. McKenna and R. J. Gaylord, Polymer 29, 2027 (1988).

64 A. Bonfanti, J. L. Kaplan, G. Charras, and A. Kabla, Soft Matter 16, 6002 (2020).

65 M. Rubinstein and S. P. Obukhov, Macromolecules 26, 1740 (1993).

66 N. Galvanetto, M. T. Ivanović, S. A. Del Grosso, A. Chowdhury, A. Sottini, D. Nettels, R. B. Best, and B. Schuler, Proceedings of the National Academy of Sciences 122 (2025), 10.1073/pnas.2424135122.

67 F. Puosi and D. Leporini, The Journal of Physical Chemistry B 115, 14046 (2011), pMID: 21793599, 10.1021/jp203659r.

68 G. Parisi, P. Urbani, and F. Zamponi, Theory of Simple Glasses: Exact Solutions in Infinite Dimensions (Cambridge University Press, 2020).

69 J. C. Phillips, Reports on Progress in Physics 59, 1133 (1996).

70 F. Arceri, F. P. Landes, L. Berthier, and G. Biroli, “Glasses and aging, a statistical mechanics perspective on,” in Statistical and Nonlinear Physics, edited by B. Chakraborty (Springer US, New York, NY, 2022) pp. 229–296.

71 E. R. Duering, K. Kremer, and G. S. Grest, The Journal of Chemical Physics 101, 8169 (1994), https://pubs.aip.org/aip/jcp/article-pdf/101/9/8169/19259761/81691online.pdf.

72 J. Li, T. Ngai, and C. Wu, Polymer Journal 42, 609–625 (2010).

73 M. Bohdan, J. Sprakel, and J. van der Gucht, Phys. Rev. E 94, 032507 (2016).

74 C. M. Fare, A. Villani, L. E. Drake, and J. Shorter, Open Biology 11, 210137 https://royalsocietypublishing.org/rsob/article-pdf/doi/10.1098/rsob.210137/935015/rsob.210137.pdf. (2021),

75 F. Dar, S. R. Cohen, D. M. Mitrea, A. H. Phillips, G. Nagy, W. C. Leite, C. B. Stanley, J.-M. Choi, R. W. Kriwacki, and R. V. Pappu, Nature Communications 15 (2024), 10.1038/s41467-024-47602-z.

76 L. Zhou, L. Zhu, C. Wang, T. Xu, J. Wang, B. Zhang, X. Zhang, and H. Wang, Nature Communications 16 (2025), 10.1038/s41467-025-58060-6.

77 L. Larini, A. Ottochian, C. De Michele, and D. Leporini, Nature Physics 4, 42–45 (2007).

78 P. J. Lu, E. Zaccarelli, F. Ciulla, A. B. Schofield, F. Sciortino, and D. A. Weitz, Nature 453, 499–503 (2008).

79 J. Wang, D. S. Devarajan, A. Nikoubashman, and J. Mittal, ACS Macro Letters 12, 1472 (2023), pMID: 37856873.

80 D. J. Bauer and A. Nikoubashman, Nature Communications 15 (2024), 10.1038/s41467-024-53575-w.

81 D. Tan, D. Aierken, P. L. Garcia, and J. A. Joseph, Soft Matter, (2025).

82 P. J. Flory, Principles of Polymer Chemistry (Cornell University Press, Ithaca, NY, 1953).

83 M. Doi and S. F. Edwards, The theory of polymer dynamics, International series of monographs on physics (Oxford Univ. Press, Oxford, 1986).

84 W. Ma, G. Zhen, W. Xie, and C. Mayr, eLife 10, e64252 (2021).

85 R. Everaers, S. K. Sukumaran, G. S. Grest, C. Svaneborg, A. Sivasubramanian, and K. Kremer, Science 303, 823 (2004), https://www.science.org/doi/pdf/10.1126/science.1091215.

86 H.-P. Hsu and K. Kremer, Macromolecules 52, 6756 (2019), pMID: 31534275.

87 M. Kröger, J. D. Dietz, R. S. Hoy, and C. Luap, Computer Physics Communications 283, 108567 (2023).

88 M. Rubinstein and A. N. Semenov, Macromolecules 34, 1058 (2001).

89 M. Rubinstein and S. Panyukov, Macromolecules 35, 6670 (2002).

90 S. Rouskin, M. Zubradt, S. Washietl, M. Kellis, and J. S. Weissman, Nature 505, 701–705 (2013).

91 K. A. Leamy, S. M. Assmann, D. H. Mathews, and P. C. Bevilacqua, Quarterly Reviews of Biophysics 49 (2016), 10.1017/s003358351600007x.

92 G. M. Wadsworth, W. J. Zahurancik, X. Zeng, P. Pullara, L. B. Lai, V. Sidharthan, R. V. Pappu, V. Gopalan, and P. R. Banerjee, Nature Chemistry 15, 1693–1704 (2023).

93 D. Aierken and J. A. Joseph, Journal of Chemical Theory and Computation 20, 10209 (2024).

94 K. Muthukumar, D. S. Devarajan, Y. C. Kim, and J. Mittal, Proceedings of the National Academy of Sciences 123, e2518384122 (2026), https://www.pnas.org/doi/pdf/10.1073/pnas.2518384122.

95 C. Roden and A. S. Gladfelter, Nature Reviews Molecular Cell Biology 22, 183–195 (2020).

96 S. Khodadadi and A. P. Sokolov, Soft Matter 11, 4984 (2015).

97 F. Bomboi, F. Romano, M. Leo, J. Fernandez-Castanon, R. Cerbino, T. Bellini, F. Bordi, P. Filetici, and F. Sciortino, Nature Communications 7 (2016), 10.1038/ncomms13191.

98 E. R. Weeks, J. C. Crocker, A. C. Levitt, A. Schofield, and D. A. Weitz, Science 287, 627 (2000), https://www.science.org/doi/pdf/10.1126/science.287.5453.627.99

99 D. Thirumalai, V. Ashwin, and J. Bhattacharjee, Phys. Rev. Lett. 77, 5385 (1996).

100 H. Tanaka, Journal of Physics: Condensed Matter 12, R207 (2000).

101 H. Maity, H. T. Nguyen, N. Hori, and D. Thirumalai, Proceedings of the National Academy of Sciences 120, e2301409120 (2023).

102 H. Maity, H. T. Nguyen, N. Hori, and D. Thirumalai, The Journal of Physical Chemistry Letters 15, 3820 (2024), https://pubs.acs.org/jpclcd/article-pdf/15/14/3820/3372530/jz3c03553.pdf.

103 S. Tian, H. Nguyen, Z. Ye, S. Rouskin, D. Thirumalai, and T. Trcek, Nature Communications 16, 8135 (2025).

104 P. G. de Gennes, The Journal of Chemical Physics 55, 572 (1971), https://pubs.aip.org/aip/jcp/article-pdf/55/2/572/18874124/5721online.pdf.

105 N. A. Erkamp, M. Farag, Y. Qiu, D. Qian, T. Sneideris, T. Wu, T. J. Welsh, H. Ausserwöger, T. J. Krug, G. Chauhan, D. A. Weitz, M. D. Lew, T. P. J. Knowles, and R. V. Pappu, Nature Communications 16 (2025), 10.1038/s41467-025-58736-z.

106 J. A. Riback, J. M. Eeftens, D. S. Lee, S. A. Quinodoz, A. Donlic, N. Orlovsky, L. Wiesner, L. Beckers, L. A. Becker, A. R. Strom, U. Rana, M. Tolbert, B. W. Purse, R. Kleiner, R. Kriwacki, and C. P. Brangwynne, Molecular Cell 83, 3095 (2023).

107 S. Jain, J. Wheeler, R. Walters, A. Agrawal, A. Barsic, and R. Parker, Cell 164, 487 (2016).

108 L. Emmanouilidis, E. Bartalucci, Y. Kan, M. Ijavi, M. E. Pérez, P. Afanasyev, D. Boehringer, J. Zehnder, S. H. Parekh, M. Bonn, T. C. T. Michaels, T. Wiegand, and F. H.-T. Allain, Nature Chemical Biology 20, 1044–1052 (2024).

109 C. He, C. Y. Wu, W. Li, and K. Xu, Journal of the American Chemical Society 145, 24240 (2023), pMID: 37782826, 10.1021/jacs.3c08674.

110 A. Miller, Z. Toprakcioglu, S. Qamar, P. St George-Hyslop, F. S. Ruggeri, T. P. J. Knowles, and M. Vendruscolo, Communications Chemistry 8 (2025), 10.1038/s42004-025-01659-z.

111 M. Linsenmeier, L. Faltova, C. Morelli, U. Capasso Palmiero, C. Seiffert, A. M. Küffner, D. Pinotsi, J. Zhou, R. Mezzenga, and P. Arosio, Nature Chemistry 15, 1340–1349 (2023).

112 C. Morelli, L. Faltova, U. Capasso Palmiero, K. Makasewicz, M. Papp, R. P. B. Jacquat, D. Pinotsi, and P. Arosio, Nature Chemistry 16, 1052–1061 (2024).

113 T. Das, F. K. Zaidi, M. Farag, K. M. Ruff, T. S. Mahendran, A. Singh, X. Gui, J. Messing, J. P. Taylor, P. R. Banerjee, R. V. Pappu, and T. Mittag, Molecular Cell 85, 2230 (2025).

114 A. Putnam, M. Cassani, J. Smith, and G. Seydoux, Nature Structural amp; Molecular Biology 26, 220–226 (2019).

115 J. Fei, M. Jadaliha, T. S. Harmon, I. T. S. Li, B. Hua, Q. Hao, A. S. Holehouse, M. Reyer, Q. Sun, S. M. Freier, R. V. Pappu, K. V. Prasanth, and T. Ha, Journal of Cell Science 130, 4180 (2017), https://journals.biologists.com/jcs/article-pdf/130/24/4180/3500924/jcs206854.pdf.

116 T. Wu, M. R. King, Y. Qiu, M. Farag, R. V. Pappu, and M. D. Lew, Nature Physics 21, 778–786 (2025).

117 S. A. Quinodoz, L. Jiang, A. A. Abu-Alfa, T. J. Comi, H. Zhao, Q. Yu, L. W. Wiesner, J. F. Botello, A. Donlic, E. Soehalim, P. Bhat, C. Zorbas, L. Wacheul, A. Košmrlj, D. L. J. Lafontaine, S. Klinge, and C. P. Brangwynne, Nature 644, 557–566 (2025).

118 I. Seim, V. Zhang, A. P. Jalihal, B. M. Stormo, S. J. Cole, J. Ekena, H. T. Nguyen, D. Thirumalai, and A. S. Gladfelter, bioRxiv (2024), 10.1101/2024.12.11.627970.

119 D. Aierken, V. Zhang, R. Sealfon, J. C. Marecki, K. D. Raney, A. S. Gladfelter, J. A. Joseph, and C. A. Roden, bioRxiv (2024), 10.1101/2024.12.23.630161.

