## Supplement Information for "Molecular origins of heterogeneous aging and spatial organization in RNA condensates"

### Supplementary information for "Molecular origins of heterogeneous aging and spatial organization of RNA condensate"


#### Structure factor

The organization of the droplet may be characterized using the static structure factor, $S(q)$. The position of a chain is computed relative to the center of mass (COM) of the droplet. For each frame in the trajectory, the COM of the entire droplet, $\mathbf{R}_{\text{droplet}}$, and the COM of each individual polymer chain $k$, $\mathbf{R}_k$, are calculated as:

$$\mathbf{R}_{\text{droplet}} = \frac{1}{N_{\text{tot}}} \sum_{i=1}^{N_{\text{tot}}} \mathbf{r}_i, \qquad \mathbf{R}_k = \frac{1}{N} \sum_{j=1}^{N} \mathbf{r}_{k,j}, \tag{1}$$

where $N_{\text{tot}}$ is the total number of beads in the droplet, $N$ is the degree of polymerization (number of beads per chain, $N = 93$ for $(\text{CAG})_{31}$ and the scramble sequence), and $\mathbf{r}_{k,j}$ is the position vector of bead $j$ in chain $k$. Each chain $k$ is assigned to a radial shell $s$ based on the Euclidean distance $d_k = \|\mathbf{R}_k - \mathbf{R}_{\text{droplet}}\|$, where the shell thickness is $\Delta r = 50$ Å.

The single-chain structure factor $S_s(q)$ for a specific shell $s$ is computed by averaging over all $M_s$ chains found within the shell (Fig. S17):

$$S_s(q) = \frac{1}{M_s N} \sum_{k=1}^{M_s} \left[ \sum_{i=1}^{N} \sum_{j=1}^{N} \frac{\sin(q r_{ij}^{(k)})}{q r_{ij}^{(k)}} \right]. \quad (2)$$

The scattering vectors $q$ are logarithmically spaced in the range $q \in [0.01, 1.0]$. In the intermediate scattering regime, the single-chain structure factor exhibits a power-law decay:

$$S_s(q) \propto q^{-\alpha}, \quad (3)$$

where the exponent $\alpha$ is related to the size of the polymer coil (Fig. S18).

#### Spatially-resolved nematic order parameter

To quantify the local orientational alignment of the polymer chains within distinct regions of the droplet, we calculate a spatially-resolved nematic order parameter, $S_s$, for each radial shell $s$. First, we assign a primary orientation vector, $\mathbf{u}_k$, to each individual chain $k$. To account for the full conformational shape of the polymer rather than just its ends, $\mathbf{u}_k$ is determined by diagonalizing the chain's $3 \times 3$ gyration tensor, $\mathbf{G}_k$:

$$\mathbf{G}_k = \frac{1}{N} \sum_{i=1}^{N} (\mathbf{r}_{k,i} - \mathbf{R}_k) \otimes (\mathbf{r}_{k,i} - \mathbf{R}_k), \quad (4)$$

where $N$ is the number of beads per chain, $\mathbf{r}_{k,i}$ is the coordinate of bead $i$ on chain $k$, and $\mathbf{R}_k$ is the COM of chain $k$. The principal axis $\mathbf{u}_k$ is the eigenvector corresponding to the largest eigenvalue of $\mathbf{G}_k$.

For a given radial shell $s$ containing $M_s$ chains, the local nematic ordering tensor $\mathbf{Q}_s$ is constructed by averaging over the orientation vectors of all chains within that shell:

$$\mathbf{Q}_s = \frac{1}{M_s} \sum_{k=1}^{M_s} \left( \frac{3}{2} \mathbf{u}_k \otimes \mathbf{u}_k - \frac{1}{2} \mathbf{I} \right), \tag{5}$$

where $\mathbf{I}$ is the $3 \times 3$ identity matrix. The local nematic director, $\hat{\mathbf{n}}_s$, representing the preferred axis of macroscopic alignment for shell $s$, is extracted as the eigenvector associated with the largest positive eigenvalue of the symmetric, traceless tensor $\mathbf{Q}_s$. Finally, the scalar nematic order parameter $P_2(\cos\theta)$ for the shell $s$ is calculated by projecting each chain's principal axis onto this local director using the second Legendre polynomial:

$$P_2 \cos\theta = \frac{1}{M_s} \sum_{k=1}^{M_s} \left[ \frac{3}{2} (\mathbf{u}_k \cdot \hat{\mathbf{n}}_s)^2 - \frac{1}{2} \right]. \tag{6}$$

A value of $P_2 \cos\theta \approx 0$ indicates an isotropic, random distribution of chain orientations, whereas $P_2 \cos\theta = 1$ corresponds to perfect alignment along $\hat{\mathbf{n}}_s$.

#### Spatially-resolved gyration tensor and shape anisotropy

Diagonalization of $\mathbf{G}_k$ in Eq. 4 yields three principal eigenvalues $(\lambda_1, \lambda_2, \lambda_3)$, which represent the squares of the principal semi-axes of the equivalent ellipsoid. For each radial shell $s$ consisting of $M_s$ chains, we calculate the following ensemble-averaged descriptors based on these eigenvalues:

- The squared radius of gyration is the trace of the gyration tensor. We define the mean $R_g$ for shell $s$ as:

$$R_{g,s} = \sqrt{\frac{1}{M_s} \sum_{k=1}^{M_s} (\lambda_1^{(k)} + \lambda_2^{(k)} + \lambda_3^{(k)})}. \tag{7}$$

- To quantify the deviation from a spherically symmetric conformation, we compute

($\kappa^2 = 0$ for a perfect sphere, $\kappa^2 = 1$ for a rigid rod):

$$\langle \kappa^2 \rangle_s = \frac{1}{M_s} \sum_{k=1}^{M_s} \left[ 1 - 3 \frac{\lambda_1^{(k)}\lambda_2^{(k)} + \lambda_2^{(k)}\lambda_3^{(k)} + \lambda_3^{(k)}\lambda_1^{(k)}}{(\lambda_1^{(k)} + \lambda_2^{(k)} + \lambda_3^{(k)})^2} \right]. \tag{8}$$

To quantify the expansion or collapse kinetics of the chains within each shell, we analyze the time evolution of the mean radius of gyration, $R_{g,s}(t)$. In the intermediate time regime, the growth rate of $\kappa^2$ is modeled as a linear process. The slope is extracted using ordinary least-squares regression.

#### Goodness-of-fit

The goodness-of-fit for a model is evaluated using the coefficient,

$$R^2 = 1 - SS_{\text{res}}/SS_{\text{tot}}. \tag{9}$$

The residual sum of squares ($SS_{\text{res}}$) measures the deviation of the observed data from the fit to the model:

$$SS_{\text{res}} = \sum_{i=1}^{n} (y_i - y(\text{model})_i)^2, \tag{10}$$

where $y_i$ is the observed value of the overlap function at time window $\tau_i$, and $y(\text{model})_i$ is the corresponding value predicted by the exponential model.

The total sum of squares ($SS_{\text{tot}}$) quantifies the total variance in the observed data relative to the mean:

$$SS_{\text{tot}} = \sum_{i=1}^{n} (y_i - \bar{y})^2, \tag{11}$$

where $\bar{y} = \frac{1}{n}\sum_{i=1}^{n} y_i$ is the arithmetic mean of the observed dataset. A value of $R^2$ (Fig. S3) close to unity indicates that the fit to the model accurately predicts the behavior of the observed function.

#### Implementation of primitive path analysis / Z1+ algorithm

To quantitatively characterize the network architecture, we employ primitive path analysis (PPA) to probe entanglement topology. Due to molecular crowding, a polymer's thermal fluctuations are confined within a tube-like region, the centerline/axis of which defines the primitive path (PP).

We calculate the PP using the Z1+ algorithm:[1] with endpoints fixed, each chain is geometrically contracted to its shortest contour without crossing its neighbors. This yields a PP of straight segments connected by kinks, representing topological constraints. The number of kinks per chain, $Z$, quantifies the degree of entanglement. The entanglement length for a chain of length $N$ can be calculated as $N_e \approx N/Z$.

To assess the spatial variations in entanglement, we assign chains to concentric shells based on their COM and calculate the average entanglements per chain per frame within each shell.

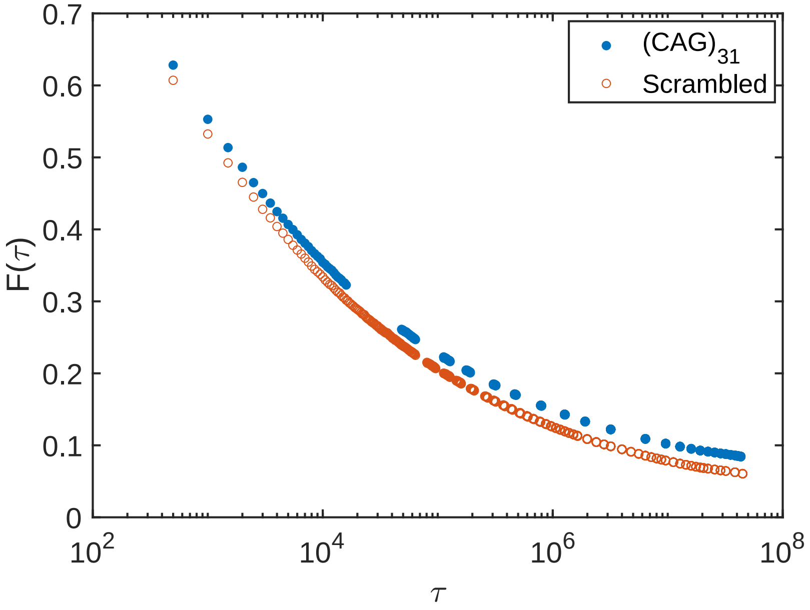


Figure S1: **Comparison of the overlap function, $F(\tau)$ (Eq. 5 in main text), for the $(CAG)_{31}$ and scrambled RNA sequences over the entire trajectory.** To isolate internal droplet dynamics, the analysis is restricted to chains that continuously reside within the condensed phase without exchanging with the dilute surrounding (30 chains for CAG repeat and 35 chains for the scrambled sequence). At all time scales, the repeat sequence (blue) has a higher $F(\tau)$ than the scrambled sequence (orange), demonstrating that the network retains greater structural memory and undergoes slower overall relaxation. Furthermore, at the longest time scales, the $CAG_{31}$ overlap function plateaus at a higher value, indicating stronger resistance to complete structural relaxation compared to the scrambled sequence. (Because the x-axis is plotted on a logarithmic scale, the initial reference point at $\tau = 0$, where $F(0) = 1$, is omitted.)

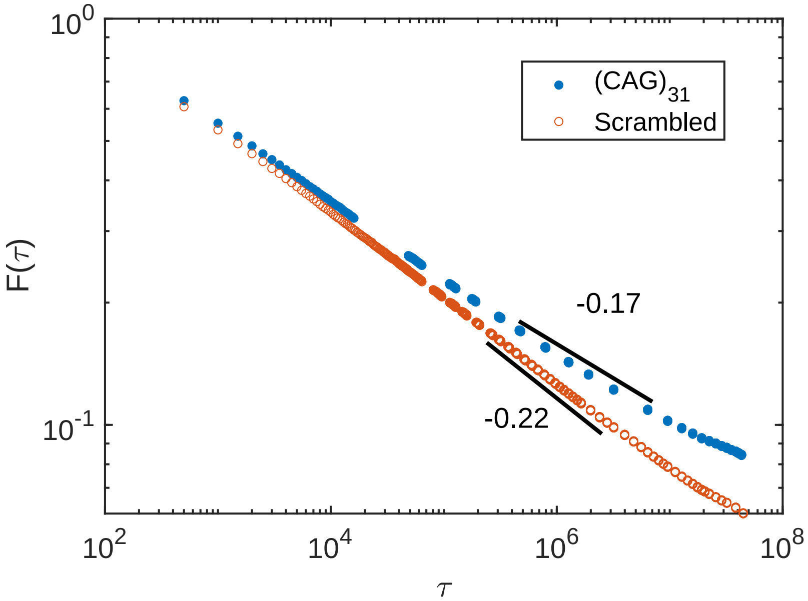


Figure S2: **Comparison of the overlap function, $F(\tau)$ (Eq. 5 in main text), for the $(CAG)_{31}$ and scrambled sequence in log-log plot.** log-log plot of overlap function decay comparison reveals underlying power law behavior of the droplet dynamics. Shallower decay ($\sim -0.17$) of $(CAG)_{31}$ sequences indicates repeat sequences retains structural memory for longer. While Scrambled sequences decay with steeper slope $\sim -0.22$ reveals they lose structural memory faster.

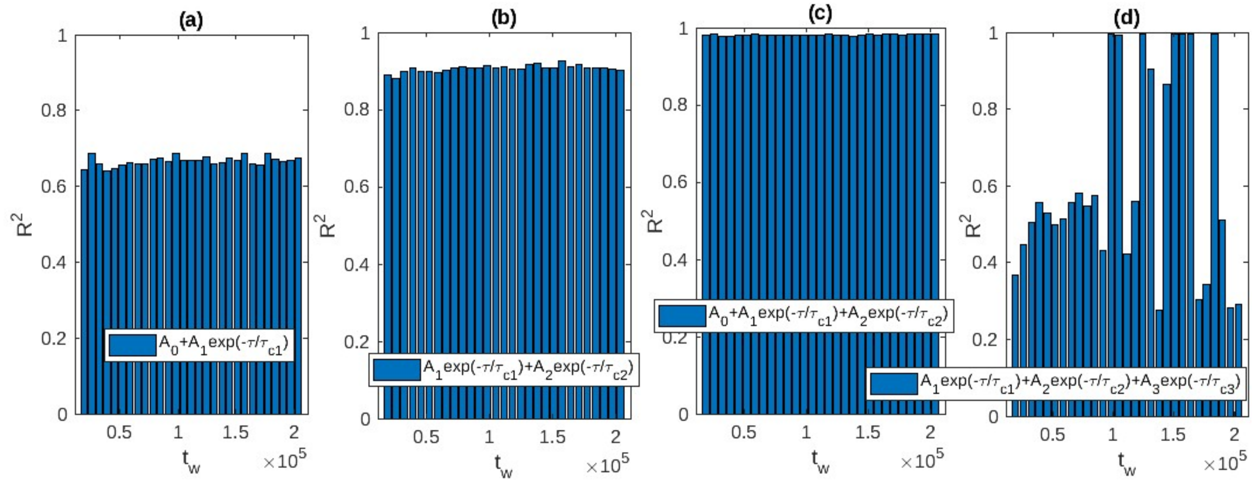


Figure S3: **Model selection for the decay of the overlap function using goodness-of-fit analysis.** The coefficient, $R^2$ (Eq. 9), is calculated for each time window ($t_w$) across four candidate models: **(a)** single exponential with offset ($A_0$), **(b)** bi-exponential without offset ($A_0$), **(c)** bi-exponential with offset ($A_0$), and **(d)** tri-exponential without offset ($A_0$). The bi-exponential model with an offset ($A_0$) (Panel c, Eq. 6 in main text) consistently yields $R^2 \approx 1$, demonstrating that it provides the optimal description of the relaxation dynamics compared to other models.

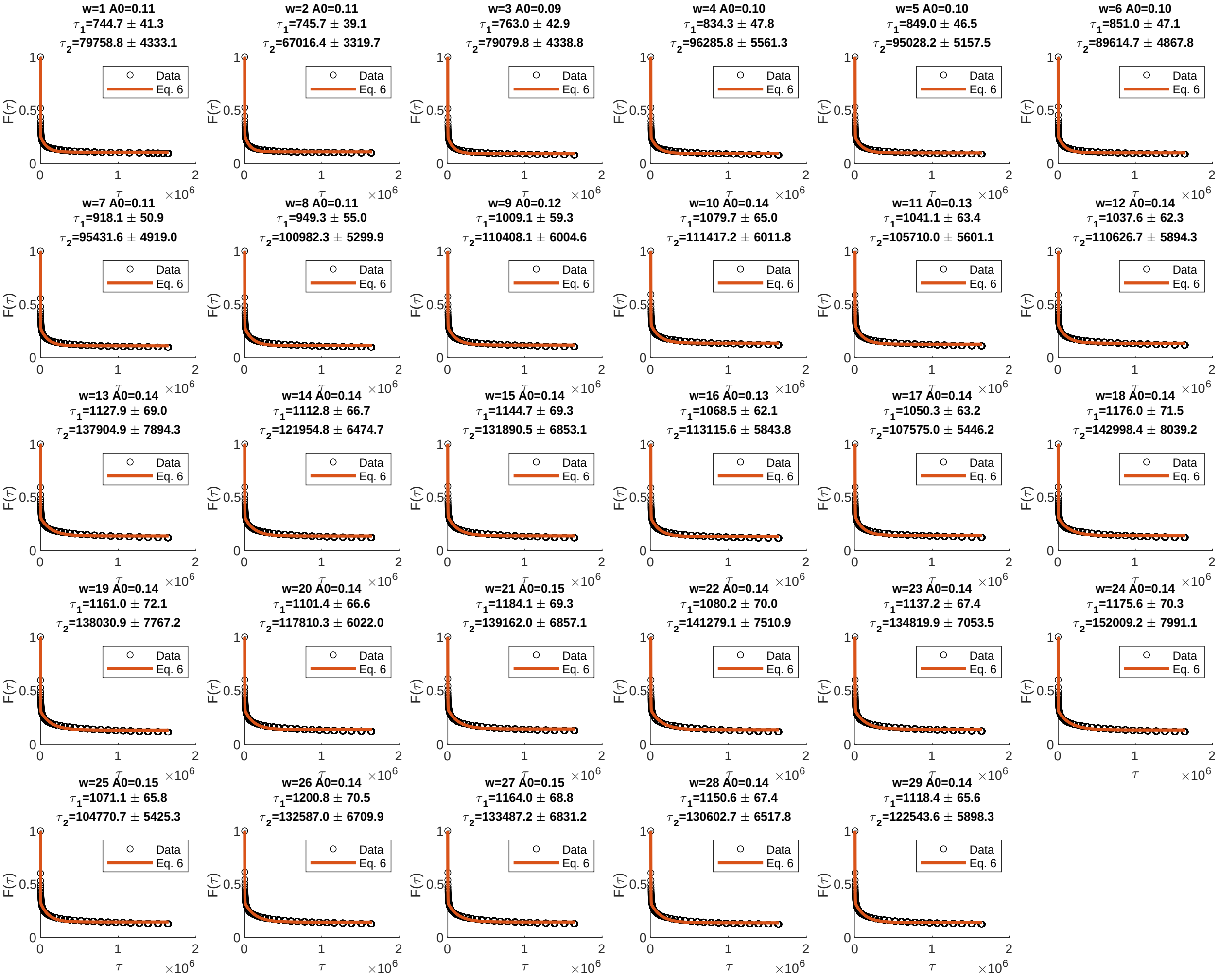


Figure S4: **Time-resolved decay of the overlap function, $F(\tau)$, across sequential time windows ($w = 1$ to 29).** In each subplot, the simulation data (open circles) are fit to the bi-exponential model described by Eq. 6 in main text (solid red line). The header of each panel displays the window index ($w$) and the extracted fitting parameters: arrested fraction ($A_0$) and characteristic relaxation times ($\tau_1, \tau_2$) along with their standard errors.

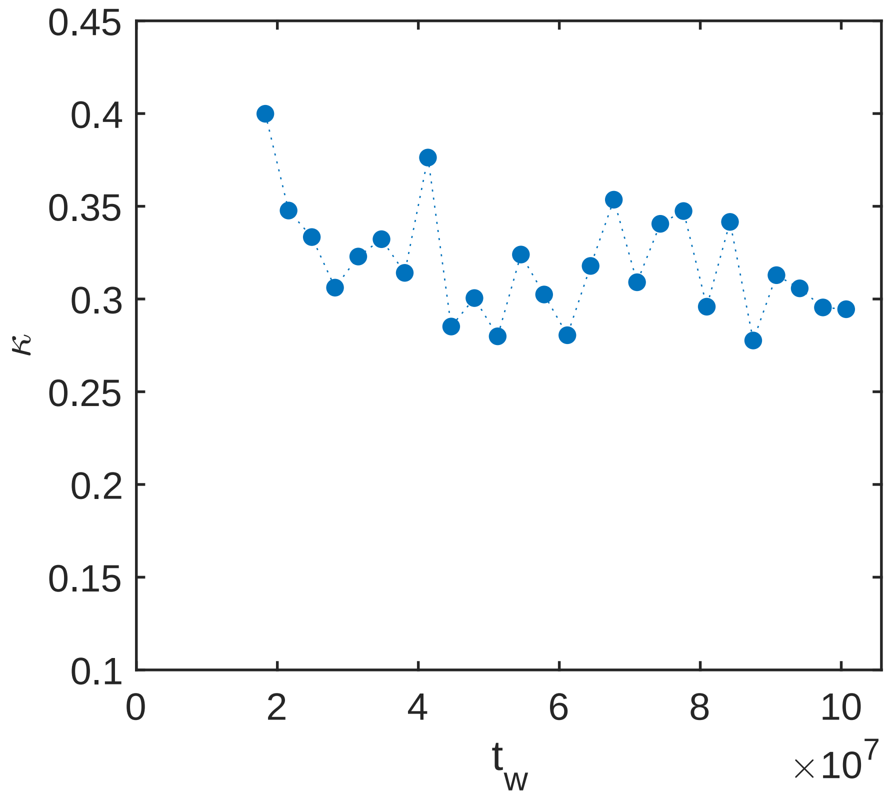


Figure S5: **Evolution of the pairwise mean square displacement (pMSD) exponent, $\kappa$, as a function of droplet age $t_w$.** $\kappa$ is derived from the scaling relation $\langle \Delta R^2_{t_w}(\tau) \rangle \sim \tau^\kappa$ (Eq. 7 in main text). The values fluctuate in the range $0.3 < \kappa < 0.4$, indicating subdiffusive dynamics across observed time windows. The mean value, $\bar{\kappa} \approx 0.35$, is used to calculate the age-dependent diffusion coefficient $D(t_w)$ (Eq. 8 in main text).

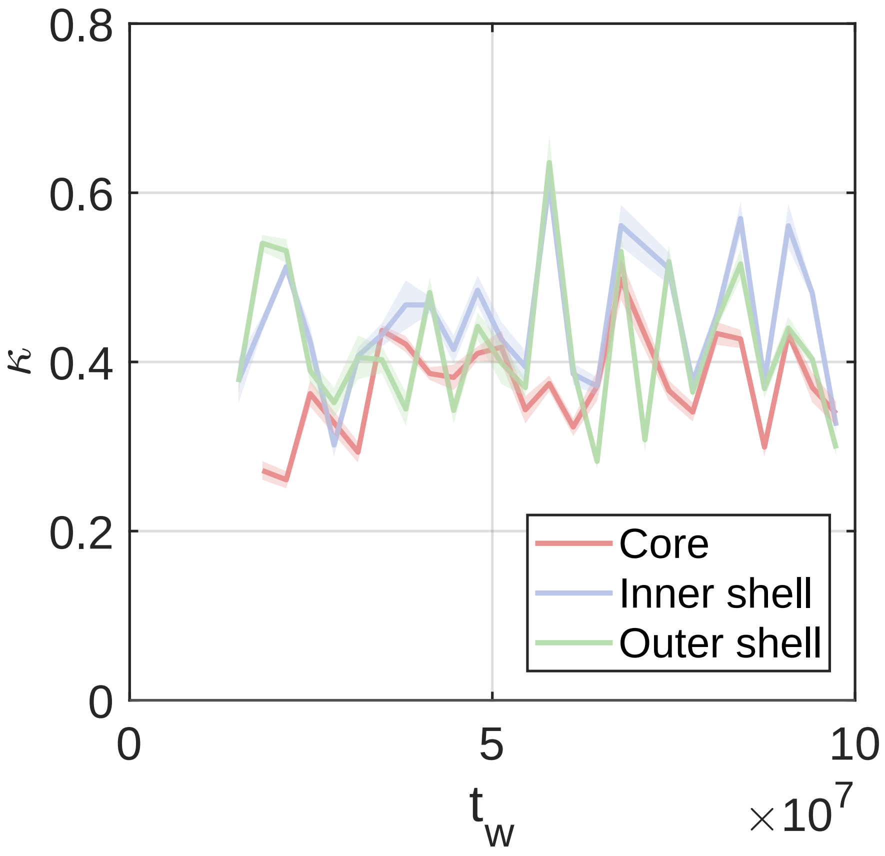


Figure S6: **MSD exponent $\kappa$ as a function of $t_w$ for different regions (core, inner shell, and outer shell) within the droplet.** The exponent values across all regions fluctuate within the range $0.3 < \kappa < 0.5$, indicating that the dynamics are consistently subdiffusive in all shells like the whole droplet. The whole droplet's mean $\bar{\kappa} \approx 0.35$, is used to calculate the age-dependent diffusion coefficient $D(t_w)$ for each shell.

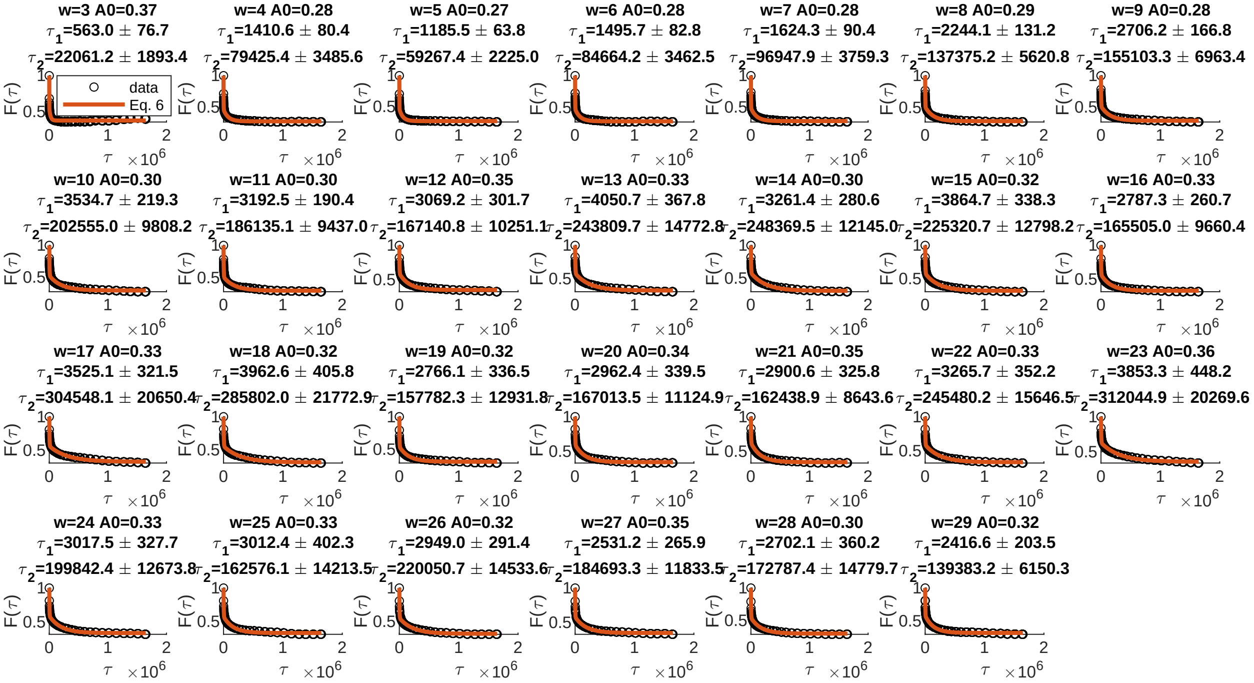


Figure S7: **Fit of the overlap function for sub-trajectories of beads located within the droplet core.**

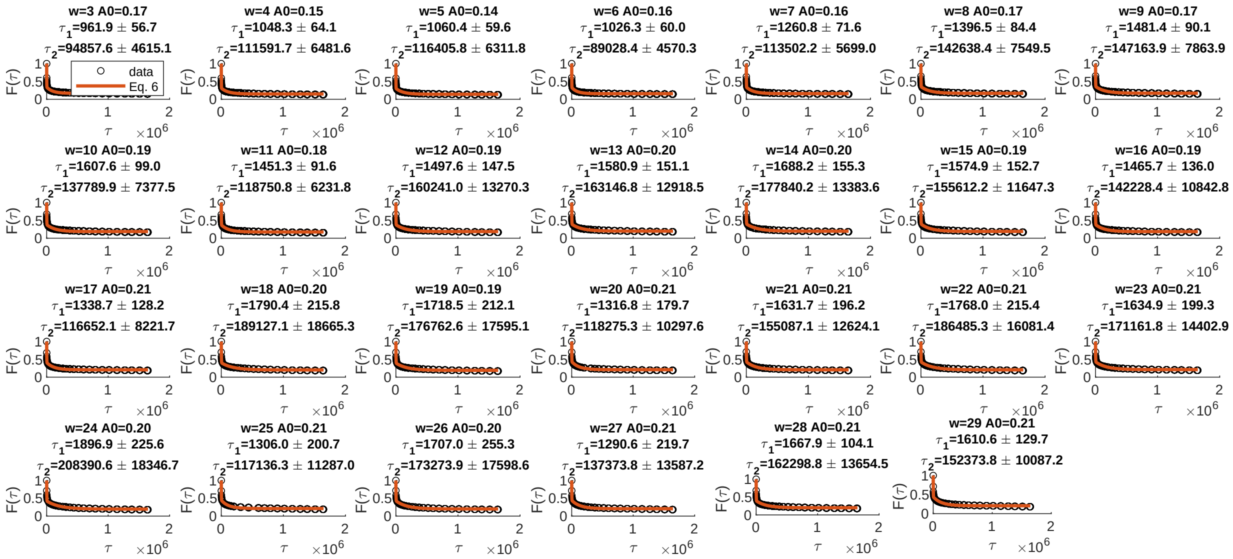

Figure S8: **Fit of the overlap function for sub-trajectories of beads located within the inner shell.**

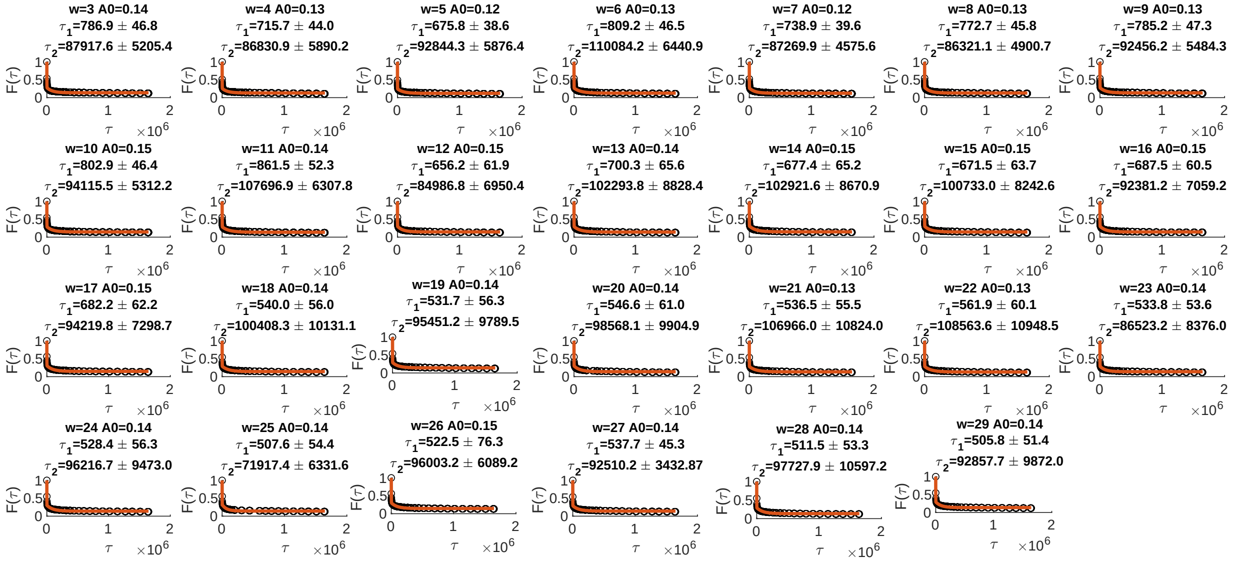


Figure S9: **Fit of the overlap function for sub-trajectories of beads located within the droplet Outer shell.**

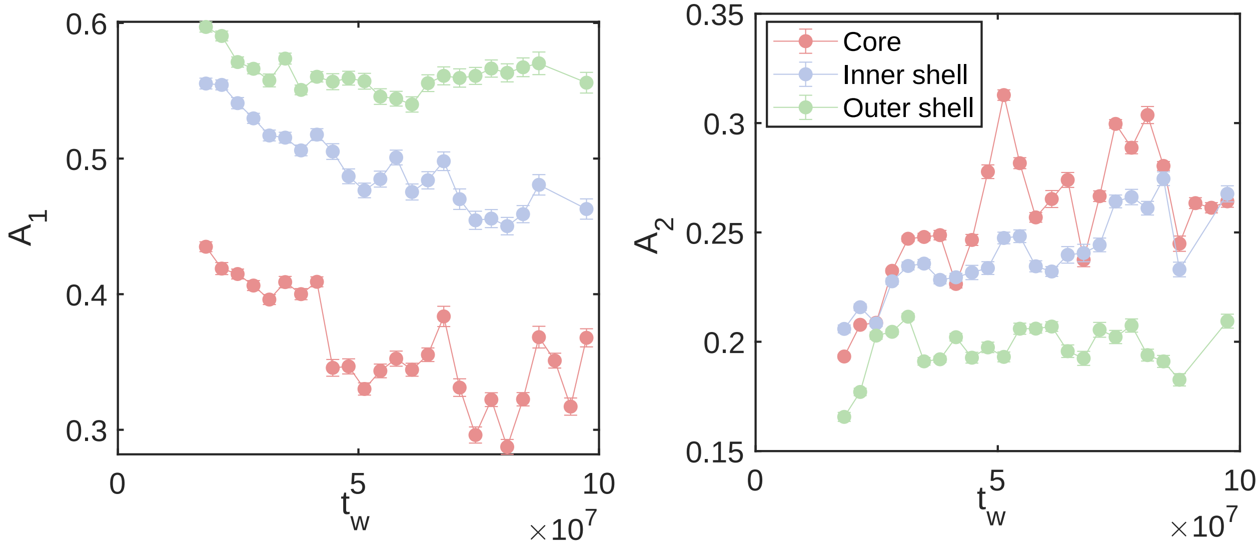


Figure S10: **Shell-wise evolution of the population fractions for the fast relaxation mode, $A_1$ (a), and the slow relaxation mode, $A_2$ (b), as the condensate ages.** The monotonic decrease of $A_1$ across all regions shows that as time increases, a progressively smaller fraction of segments relaxes via the fast mode in all the shells. In contrast, the slow-mode population $A_2$ increases in the core and inner shells, but only modestly in the outer shell.

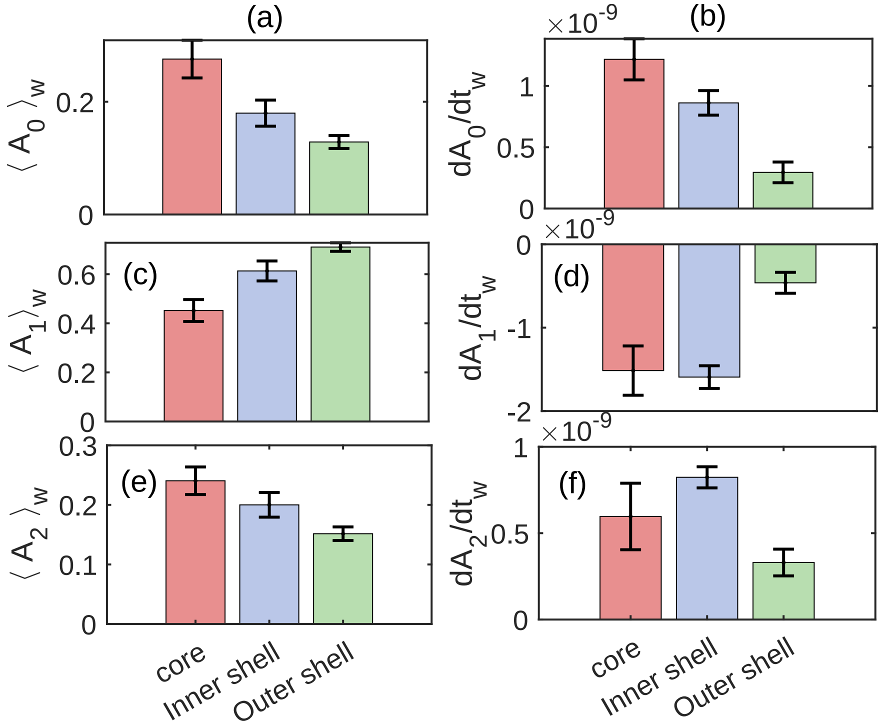


Figure S11: **Shell-dependent time-average and rate of population fractions** (a) Comparison of the mean $\langle A_0 \rangle$ over all time-windows between core, inner and outer shell. Average $A_0$ is a maximum at in the core and decreases towards the periphery. (b) The rate of change of $A_0$ is also a maximum at the core and decreases toward the surface gradually. This indicates that the early solidification is initiated at the core and spreads toward the inner and outer shells as time progresses. (c) Mean population fraction of fast relaxation $A_1$ is maximum at core, and decreases towards surface. (d) Rate of decrease of $A_1$ is largest at core and inner shell, compared to outer shell. This indicates chain segments do not relax through fast relaxation mode as the droplet ages within the interior of the droplet, compared to outer shell. (e) Mean population fraction of fast relaxation $A_2$ is maximum at core, decreases towards surface. (f) Rate of increase of $A_2$ is large in the interior of the droplet and decreases at the outer shell.

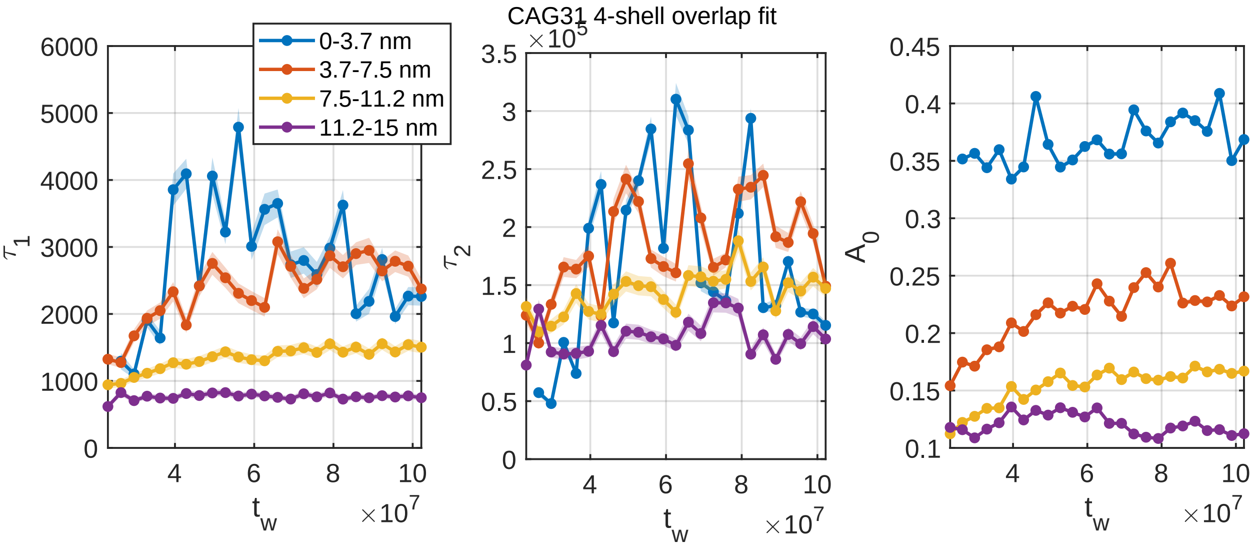


Figure S12: **Shell-resolved relaxation of a CAG31 condensate is preserved with four shells.** The pair-overlap function is decomposed into four concentric shells of $\approx 3.75$ nm thickness (0–3.7, 3.7–7.5, 7.5–11.2, and 11.2–15 nm from the droplet COM). Panels (left to right) show the fast relaxation time $\tau_1$, the slow relaxation time $\tau_2$, and the arrested fraction $A_0$ as functions of the waiting time $t_w$. Shaded bands denote the fit uncertainty. All three quantities decrease from the core to the surface at every $t_w$.

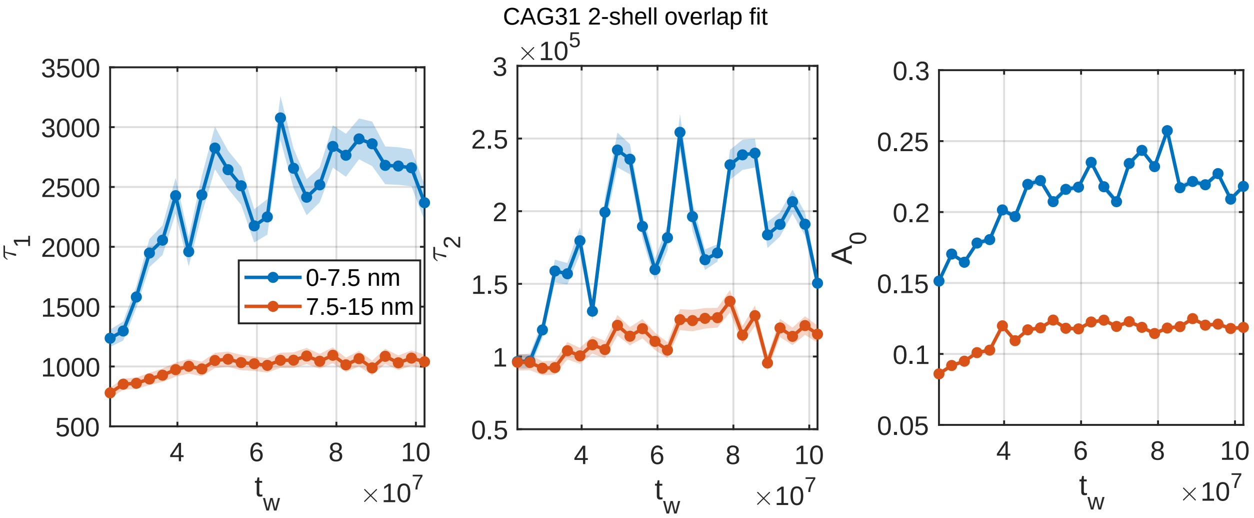


Figure S13: **The same behaviour holds with two shells.** The overlap function is coarse-grained into an interior shell (0–7.5 nm) and a surface shell (7.5–15 nm), each 7.5 nm thick, and fitted as in the four-shell case. The fast time $\tau_1$, slow time $\tau_2$, and arrested fraction $A_0$ (left to right) remain systematically larger in the interior than at the surface throughout aging. Shaded bands denote the fit uncertainty.

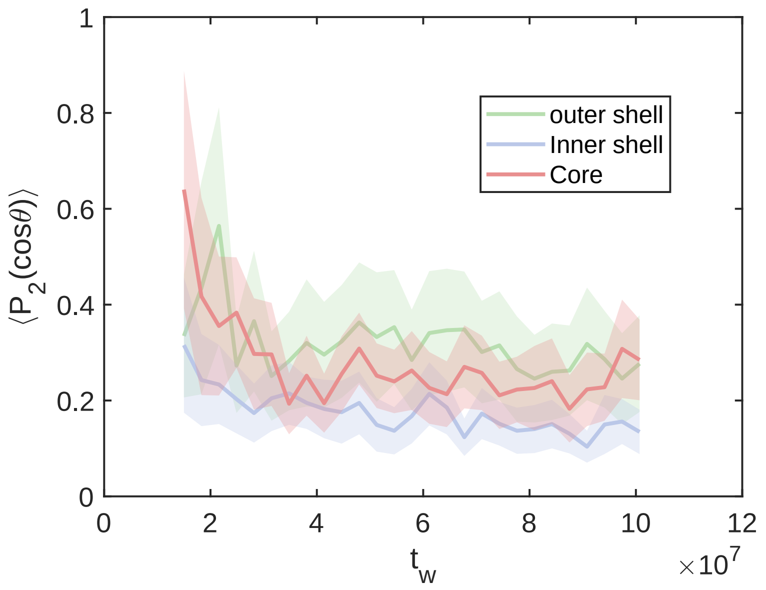


Figure S14: **Time evolution of the local nematic order parameter, $S = \langle P_2(\cos\theta)\rangle$, for each region.** The parameter quantifies the degree of orientational alignment of the polymer chains relative to the local director within each spatial region.

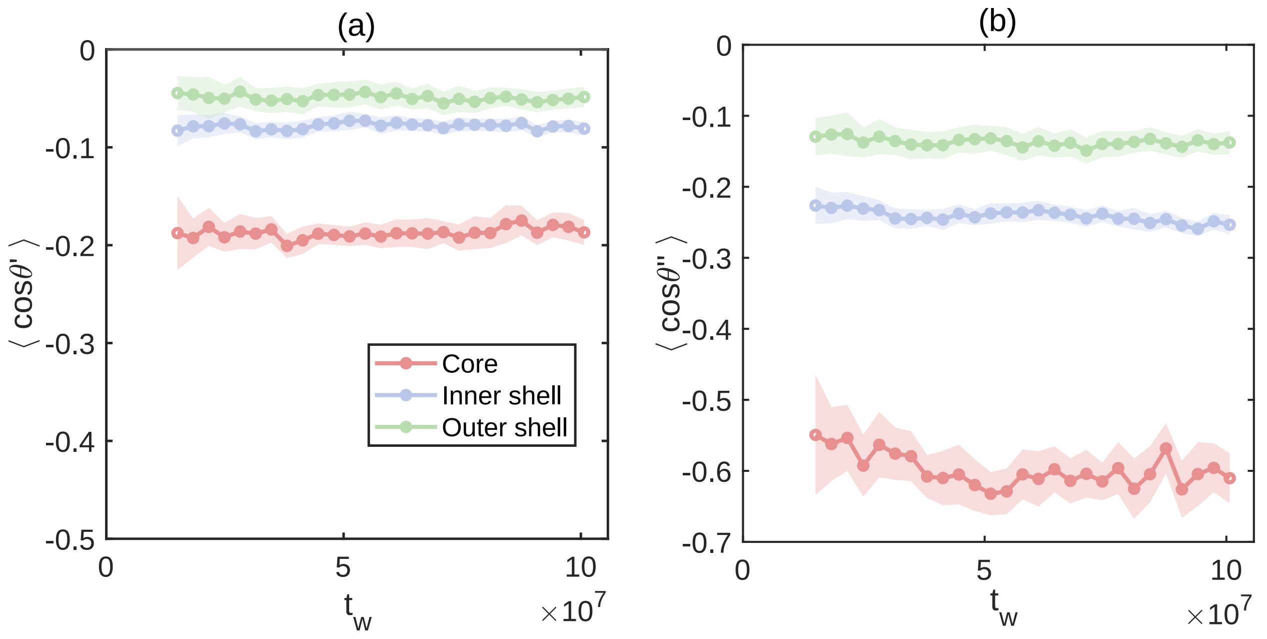


Figure S15: **Robustness of radial chain orientation across different segment lengths.** To verify that the radial alignment discussed in the main text (Fig. 3d) calculated for segment separation $\Delta i = 6$ is not an artifact of the chosen length scale, we analyze the orientation for alternative separations: **(a)** short-range ($\Delta i = 3$) and **(b)** long-range ($\Delta i = 10$). The alignment parameter is defined as the average cosine of the angle between the radial vector from the droplet center of mass ($\vec{r}_{\text{COM}\to i}$) and the segment vector ($\vec{r}_{i\to i+\Delta i}$). In both cases, the core region (red) exhibits significantly more negative values compared to the inner (blue) and outer (green) shells. This confirms that the distinct radial ordering of the polymer chains within the droplet core is a robust feature that persists across different length scales, becoming more pronounced for longer segments.

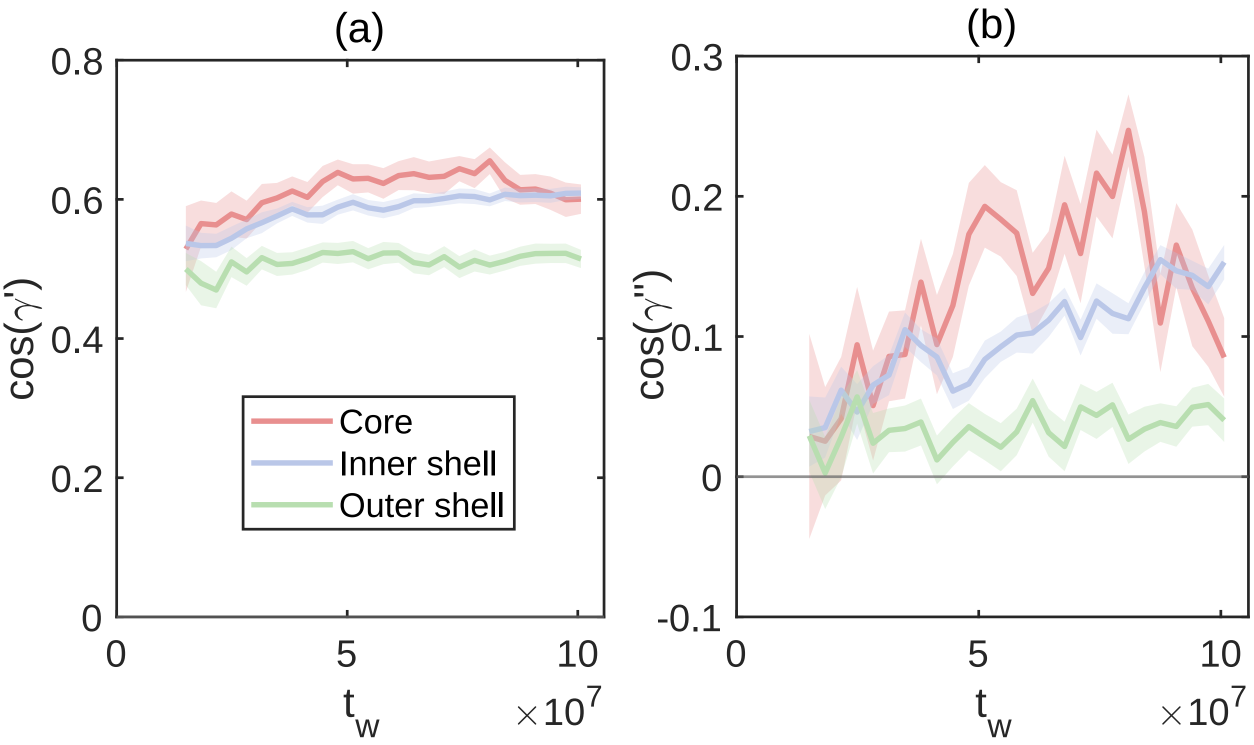


Figure S16: **Robustness of bond orientational correlation across different segment separations.** To show the robustness of the bond orientational analysis presented in the main text (based on separation $\Delta i = 6$), we compute the same quantity separated by different **(a)** $\Delta i = 3$ beads ($\langle \cos\gamma \rangle = \langle \hat{u}_i \cdot \hat{u}_{i+3} \rangle$) and **(b)** $\Delta i = 10$ beads ($\langle \cos\gamma' \rangle = \langle \hat{u}_i \cdot \hat{u}_{i+10} \rangle$). Consistent with the main text findings, the droplet core (red) maintains a moderate positive correlation.

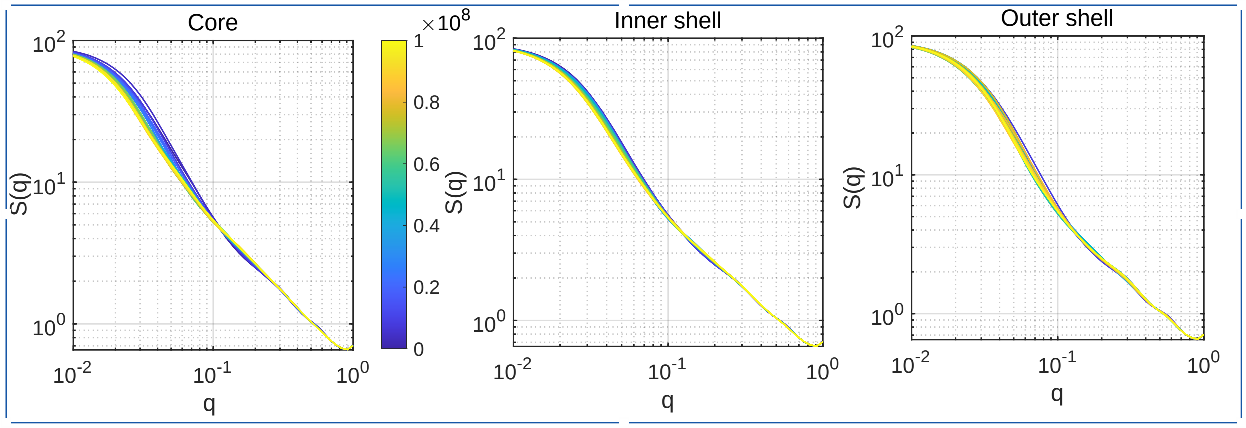


Figure S17: **Variation of single-chain static structure factor, $S(q)$, at different time windows $t_w$ across different radial shells.** The color gradient from blue to yellow indicates the progression of condensate age, $t_w$. In the intermediate scattering regime ($0.03 \leq q \leq 0.3$ Å$^{-1}$), the curves exhibit the characteristic power-law decay.

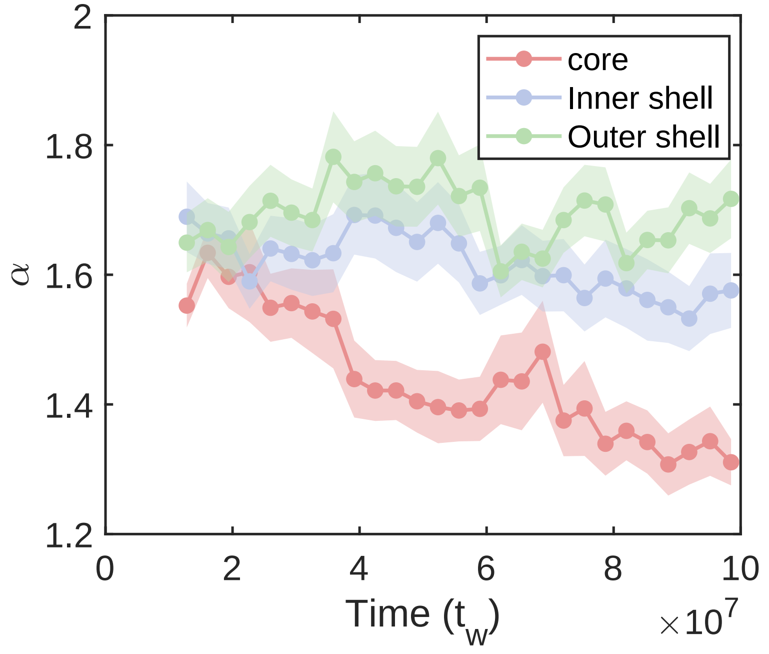


Figure S18: **Time evolution of the structural scaling exponent, $\alpha$, across different radial shells.** The exponent is extracted by fitting the intermediate scattering regime ($0.03 \leq q \leq 0.3$ Å$^{-1}$) of the single-chain static structure factor in Fig. S14 to a power-law, $S(q) \propto q^{-\alpha}$. The outer shell (green) and inner shell (blue) maintain relatively higher $\alpha$ values between 1.6 and 1.8, indicative of polymer coil conformations. In contrast, the core region (red) exhibits a distinct, monotonic decrease in $\alpha$ over time (from $\approx 1.6$ to $\approx 1.3$). This progressive reduction in $\alpha$ within the core suggests chains within the dense droplet interior gradually adopt more extended configurations as the condensate ages.

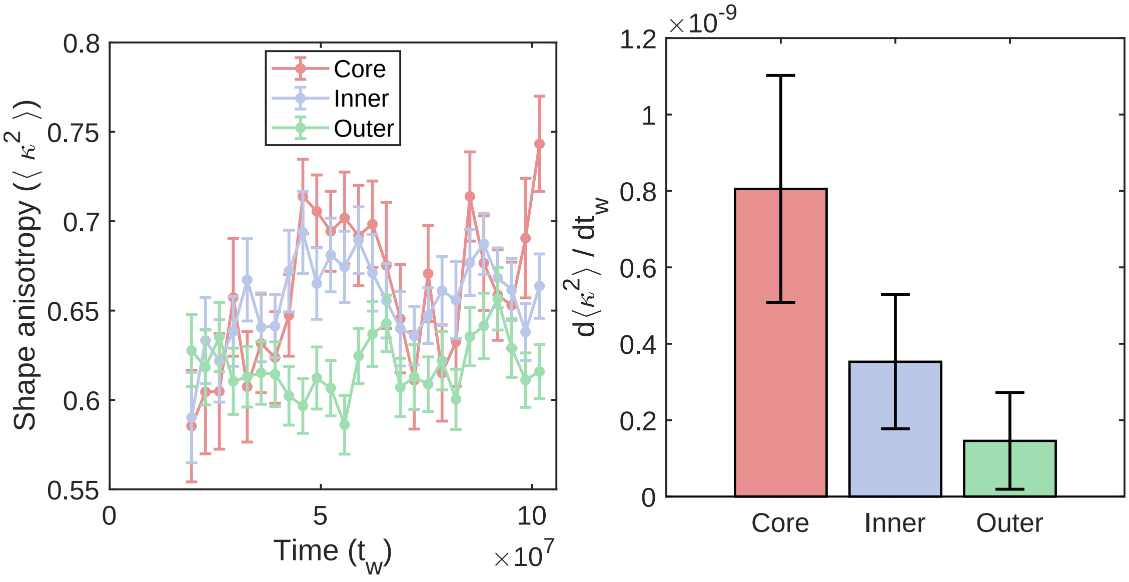


Figure S19: **Shellwise shape anisotropy variation of chains within the droplet** (a) Shellwise shape anisotropy variation with condensate age $t_w$. Chain segments within the core are more anisotropic (non-spherical) compared to those on the periphery. (b) The rate of increase of shape anisotropy is highest in the core region and decreases significantly towards the surface, where it remains nearly unchanged.

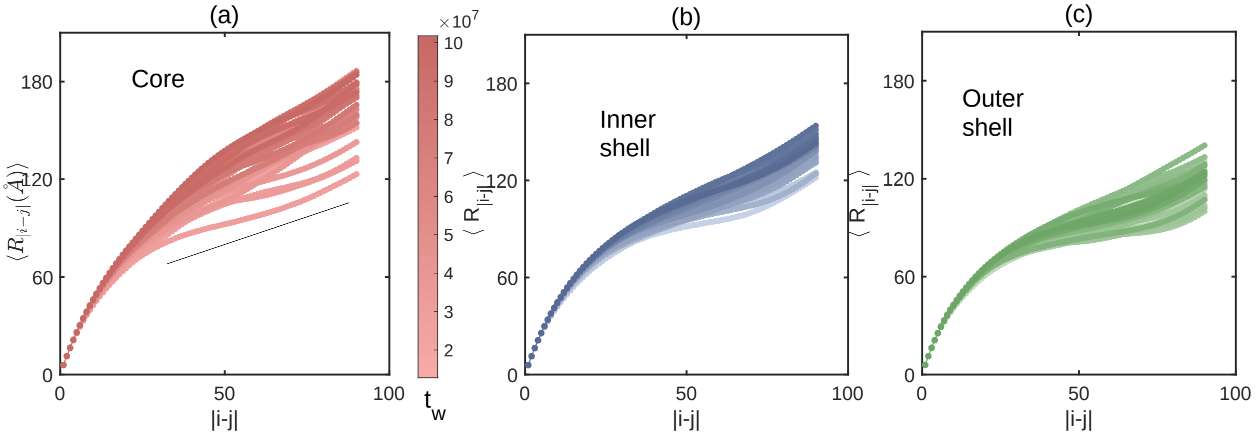


Figure S20: **Shell-resolved segmental size of droplet chains.** Shellwise variation of the mean segmental size $R_{|i-j|}$ as a function of contour separation $|i-j|$ for chains whose COM lies in the indicated shell. The color gradient (light → dark) indicates increasing $t_w$. Power-law fits $R_{|i-j|} \propto |i-j|^{\nu}$, used to extract the segmental size exponent $\nu$, are taken over the linear regime $40 < |i-j| < 90$ and reported in Fig. 3c in main text. Chains in the core are more extended and display larger $R_{|i-j|}$, while chains in the inner shell are less extended and those in the outer shell are most restricted.

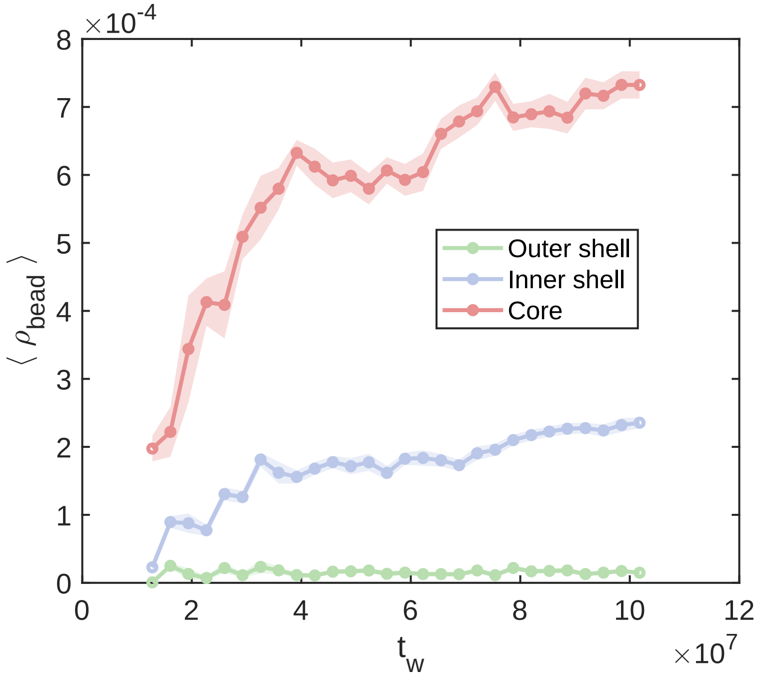


Figure S21: **Shell-resolved evolution of bead density.** Mean bead density $\langle \rho_{\text{bead}} \rangle$ in each shell is defined as $\rho_{\text{bead}} = \sum N_s/V_s$, where $N_s$ is the total number of beads in a given shell with volume $V_s$. The core density increases strongly with condensate age $t_w$ and becomes about 3–4 times higher than in the inner shell, while the outer-shell density remains low and unchanged. The corresponding growth rates of density (slopes of $\langle \rho_{\text{bead}} \rangle$ vs. $t_w$) are shown in Fig. 5b in main text.

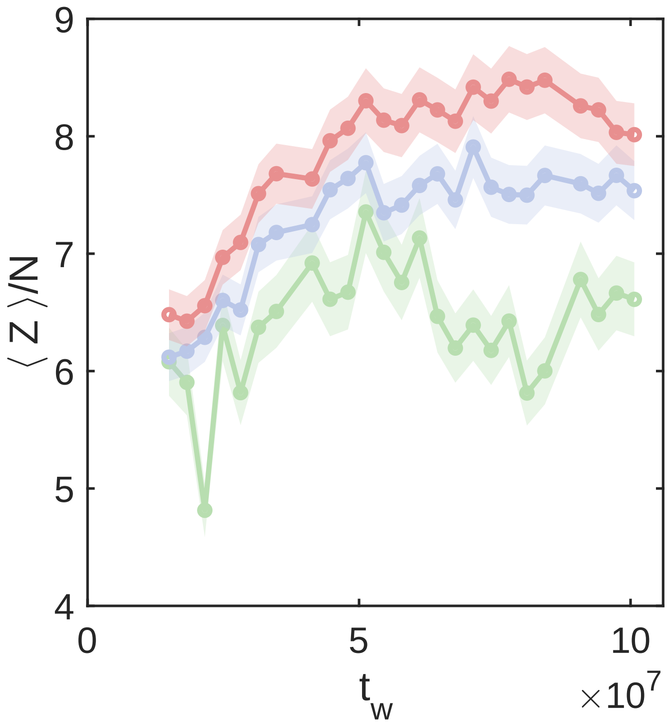


Figure S22: **Shell-resolved topological entanglements per chain, $\langle Z \rangle / N$, as the condensate ages.** The number of entanglements (kinks) is determined using the Z1+ algorithm. The core region (red) consistently exhibits a higher density of entanglements compared to the inner (blue) and outer (green) shells. Furthermore, the core shows a pronounced, progressive increase in $\langle Z \rangle / N$ over time, indicating that the polymer network within the droplet interior becomes increasingly topologically constrained as it ages. This spatial and temporal entanglement trend is in strong agreement with the spatial gradients observed in the bead and base-pair densities.

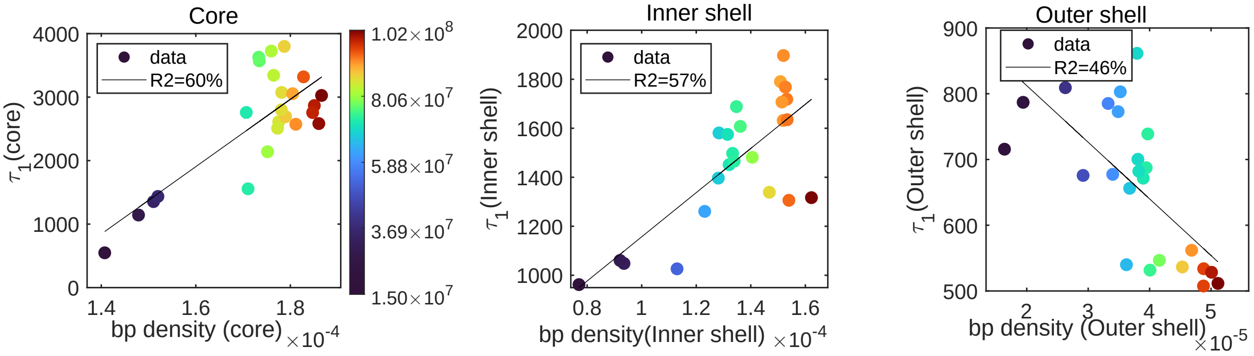


Figure S23: **Shell-wise correlation between local base-pair (bp) density and the fast relaxation timescale, $\tau_1$.** Data points (open circles) correspond to paired measurements extracted at matched time windows ($t_w$) for the **(a)** Core, **(b)** Inner shell, and **(c)** Outer shell. Solid lines represent linear regressions, with the goodness-of-fit indicated by the $R^2$ values.

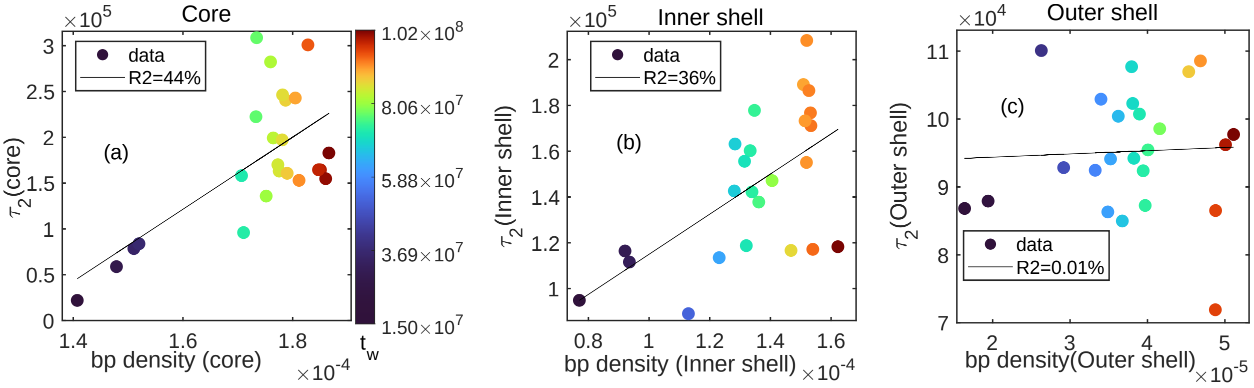


Figure S24: **Shell-wise correlation between local base-pair (bp) density and the slow relaxation timescale, $\tau_2$.** Similar to the fast relaxation mode, the core (a) and inner shell (b) display positive correlations ($R^2 = 0.44$ and 0.36), linking increased structural density to prolonged slow-mode relaxation. The outer shell (c) exhibits no such correlation ($R^2 \approx 0$).

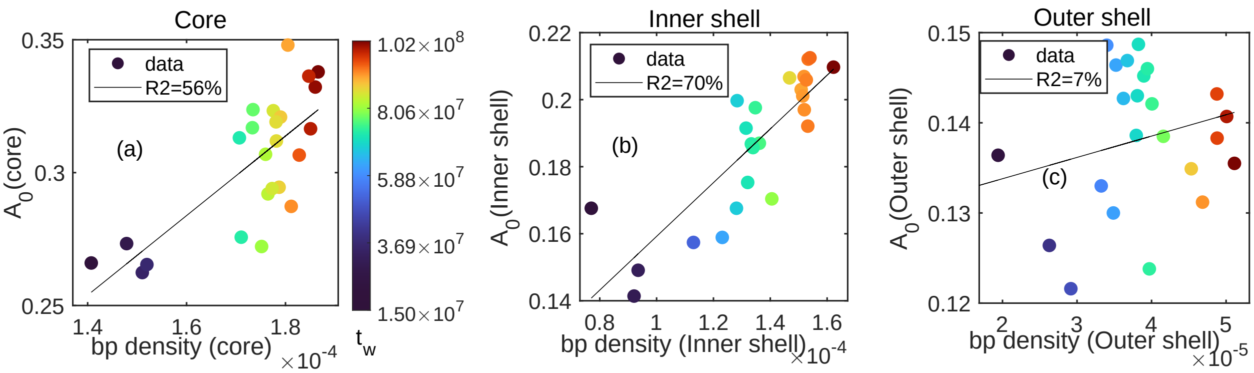


Figure S25: **Shellwise correlation between bp density and non-ergodic population fraction** $A_0$. Positive correlation is observed in the core ($R^2 = 0.56$) (a) and inner shell ($R^2 = 0.70$) (b), demonstrating that increased packing density directly enhances the solid-like, arrested fraction of the network. $A_0$ shows little correlation with bp density in outer shell with $R^2 = 7\%$ (c).

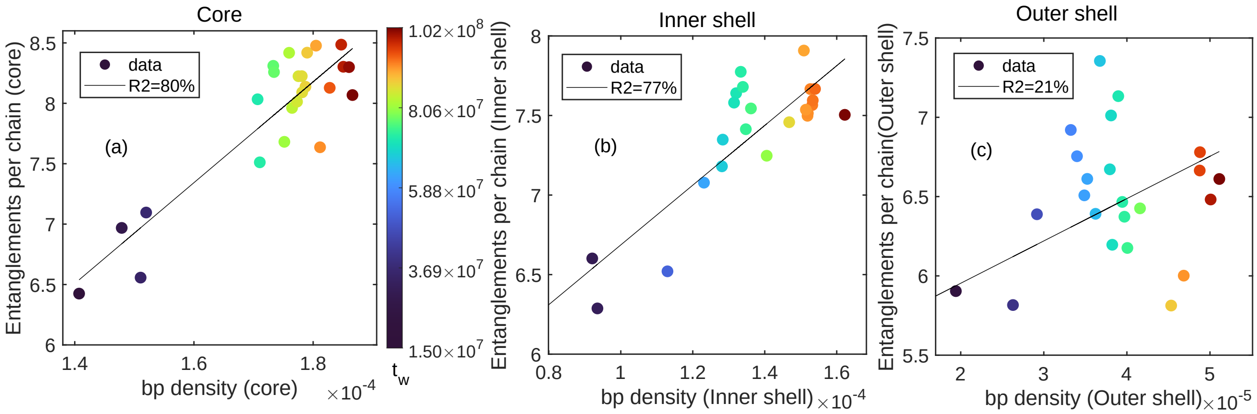


Figure S26: **Shell-wise correlation between local base-pair (bp) density and topological entanglements per chain, $\langle Z \rangle / N$.** The core (a) and inner shell (b) exhibit highly positive correlations ($R^2 = 0.80$ and $0.77$), providing direct evidence that spatial density gradients are tightly coupled to the local degree of topological cross-linking. The outer shell (c) exhibits a significantly weaker dependence ($R^2 = 0.21$).

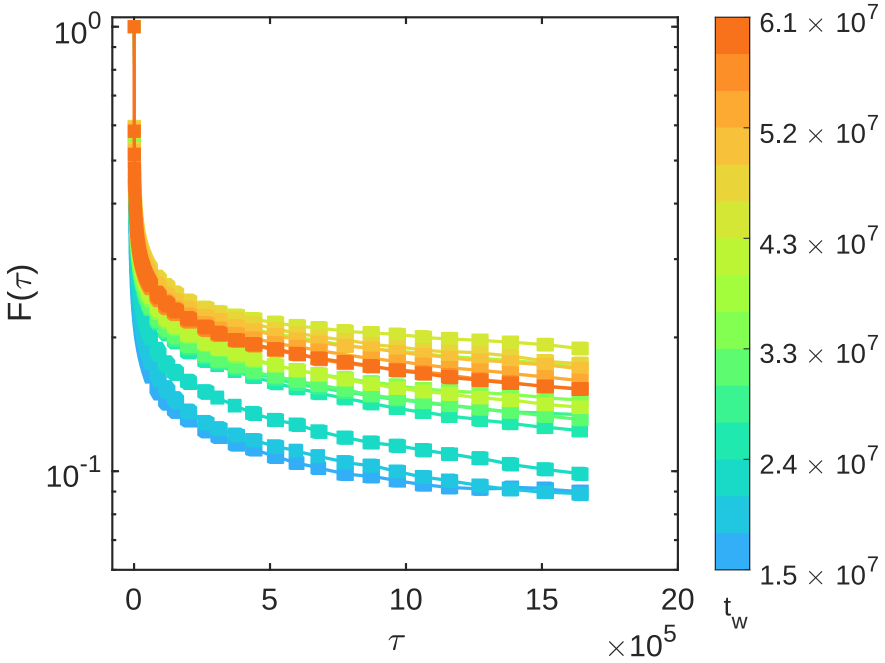


Figure S27: **Time evolution of the overlap function, $F(\tau)$, for the scrambled RNA sequence.** The decay of the overlap function is plotted against observation time $\tau$. The color gradient from blue to orange corresponds to increasing condensate age, $t_w$.

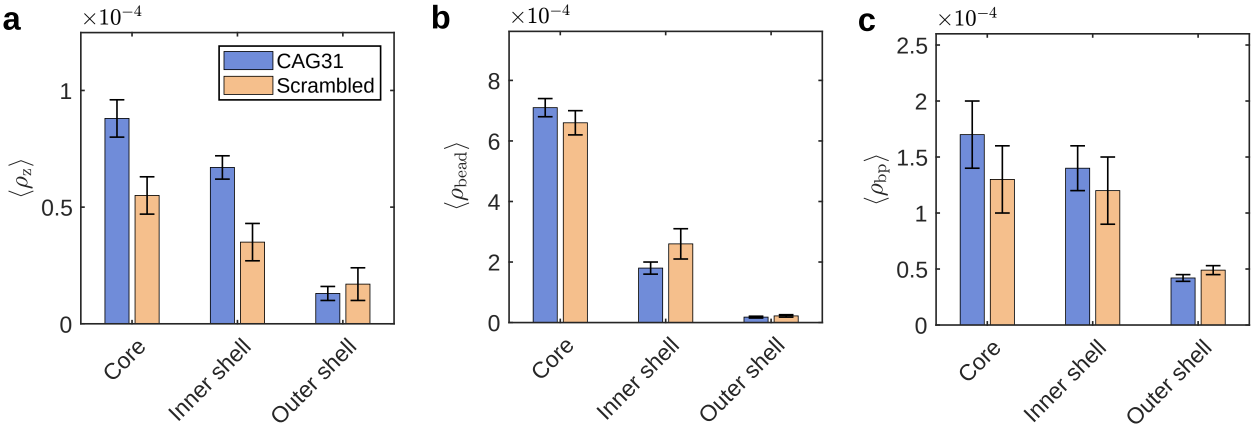


Figure S28: Entanglement density, bead density and base-pair density comparison between $(CAG)_{31}$ and scrambled sequences when both of the condensates are matured at the end of our simulations.

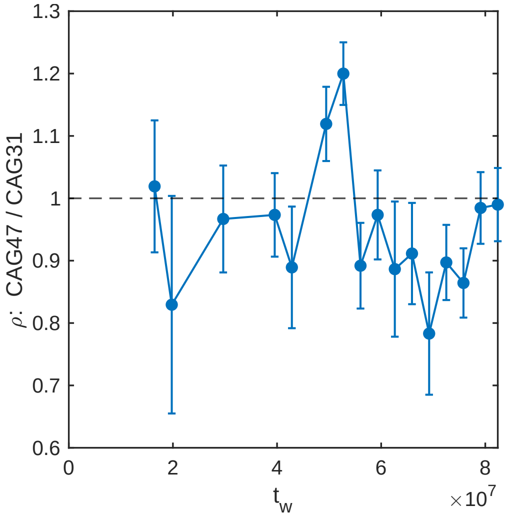


Figure S29: **Bead density ratio between $CAG_{47}$ and $CAG_{31}$ condensates**: The ratio stays close to unity throughout aging, showing that the two condensates reach comparable overall density.

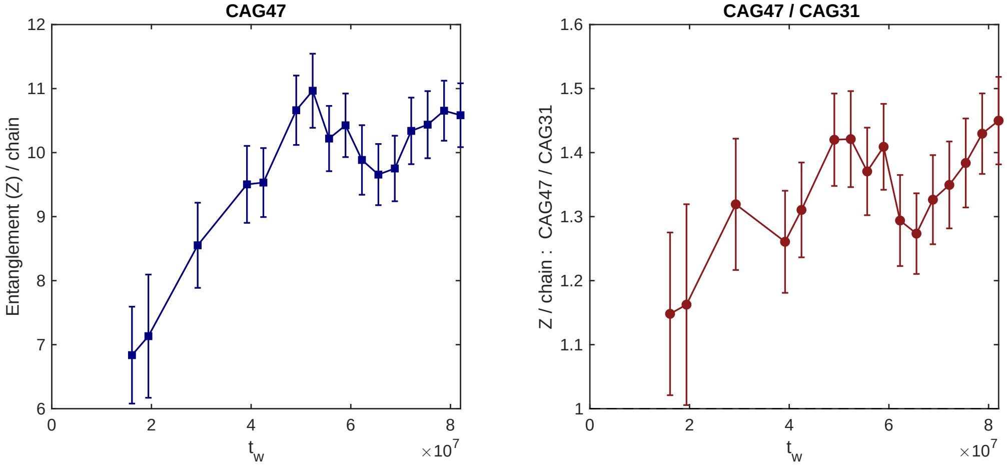


Figure S30: **Entanglements per chain ratio between $CAG_{47}$ and $CAG_{31}$**: The ratio remains 30–40% above unity throughout aging, showing that entanglement tracks chain length independently of density.

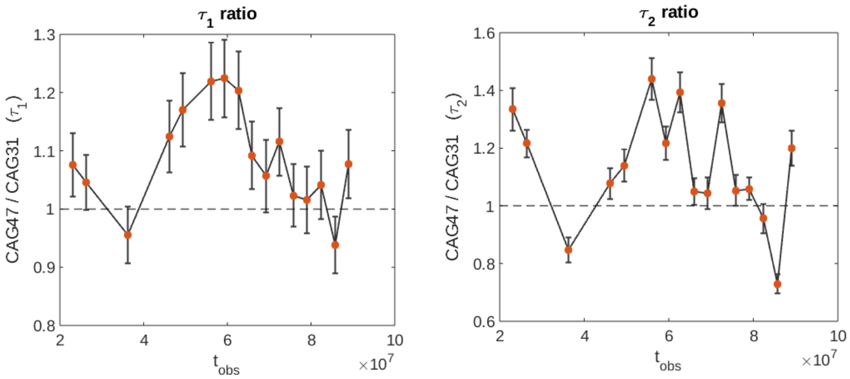


Figure S31: **Ratio of relaxation timescales ($\tau_1$ and $\tau_2$) between $CAG_{47}$ and $CAG_{31}$**: Both ratios stay above unity throughout aging, showing slower relaxation in the longer $CAG_{47}$ chains.

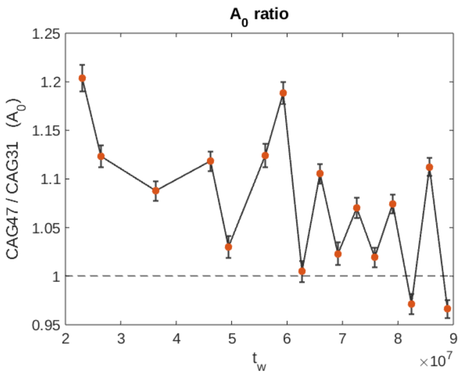


Figure S32: **Ratio of arrested fraction $A_0$ between $CAG_{47}$ and $CAG_{31}$**: The ratio remains above unity, indicating a larger non-relaxing (arrested) population in $CAG_{47}$ relative to $CAG_{31}$.

### References

(1) Kröger, M.; Dietz, J. D.; Hoy, R. S.; Luap, C. The Z1+ package: Shortest multiple disconnected path for the analysis of entanglements in macromolecular systems. *Computer Physics Communications* **2023**, *283*, 108567.
